# Comprehensive structure-function analysis reveals gain- and loss-of-function mechanisms impacting oncogenic KRAS activity

**DOI:** 10.1101/2024.10.22.618529

**Authors:** Jason J. Kwon, Julien Dilly, Shengwu Liu, Eejung Kim, Yuemin Bian, Srisathiyanarayanan Dharmaiah, Timothy H. Tran, Kevin S. Kapner, Seav Huong Ly, Xiaoping Yang, Dana Rabara, Timothy J. Waybright, Andrew O. Giacomelli, Andrew L. Hong, Sean Misek, Belinda Wang, Arvind Ravi, John G. Doench, Rameen Beroukhim, Christopher T. Lemke, Kevin M. Haigis, Dominic Esposito, David E. Root, Dwight V. Nissley, Andrew G. Stephen, Frank McCormick, Dhirendra K. Simanshu, William C. Hahn, Andrew J. Aguirre

**Affiliations:** Department of Medical Oncology, Dana Farber Cancer Institute, Boston, MA, 02115, USA; Harvard Medical School, Boston, MA, 02115, USA; Broad Institute of MIT and Harvard, Cambridge, MA, 02142, USA; NCI RAS Initiative, Cancer Research Technology Program, Frederick National Laboratory for Cancer Research, Frederick, MD, USA; Humber Polytechnic, Toronto, ON, Canada; Department of Pediatrics, Emory University School of Medicine and Children’s Healthcare of Atlanta, Atlanta, GA, USA; University of California, San Francisco Helen Diller Family Comprehensive Cancer Center, University of California, San Francisco, CA, USA

**Keywords:** KRAS, cell transformation, structure-function, deep mutational scanning, cancers

## Abstract

To dissect variant-function relationships in the KRAS oncoprotein, we performed deep mutational scanning (DMS) screens for both wild-type and KRAS^G12D^ mutant alleles. We defined the spectrum of oncogenic potential for nearly all possible *KRAS* variants, identifying several novel transforming alleles and elucidating a model to describe the frequency of *KRAS* mutations in human cancer as a function of transforming potential, mutational probability, and tissue-specific mutational signatures. Biochemical and structural analyses of variants identified in a KRAS^G12D^ second-site suppressor DMS screen revealed that attenuation of oncogenic KRAS can be mediated by protein instability and conformational rigidity, resulting in reduced binding affinity to effector proteins, such as RAF and PI3-kinases, or reduced SOS-mediated nucleotide exchange activity. These studies define the landscape of single amino acid alterations that modulate the function of KRAS, providing a resource for the clinical interpretation of KRAS variants and elucidating mechanisms of oncogenic KRAS inactivation for therapeutic exploitation.

## Introduction

KRAS is a monomeric GTPase that functions as a molecular switch to control cell proliferation, survival, and differentiation*. KRAS* is the most commonly mutated oncogene in cancer, especially in adenocarcinomas of the pancreas, colon, and lung^1^. The majority of *KRAS* mutations occur at hotspot codons 12, 13, and 61^1^. Missense mutations at these positions interfere with GTP hydrolysis and/or nucleotide exchange, and thus increase the steady state level of KRAS-GTP. The resulting constitutive activation of oncogenic *KRAS*, combined with concurrent inactivation of tumor suppressor genes, promotes tumor formation in mouse models of pancreatic, lung, and colon cancers, and other *in vitro* and *in vivo* transformation models^2–4^. The proliferation of cancer cell lines and genetically engineered mouse tumors that harbor oncogenic *KRAS* is dependent on continued KRAS expression^5,6^.

Human tumors are now routinely sequenced, resulting in the identification of several rare *KRAS* variants with unknown functions. The Catalogue Of Somatic Mutations In Cancers (COSMIC) lists more than 240 variants of KRAS identified in human cancer, with about half of these variants reported to be private mutations in individual patients^7^. Functional investigation of these variants is critical to better understand their oncogenic potential. Moreover, specific *KRAS* mutations have been associated with distinct clinical prognoses and varying responses to chemotherapy and may also serve as a predictive marker for the effectiveness of certain targeted therapies^8^. For example, metastatic colorectal cancers that harbor oncogenic *KRAS* mutations do not benefit from treatment with monoclonal antibodies targeting EGFR^8,9^.

KRAS^G12C^ became the first KRAS mutant that can be directly targeted through the covalent binding of small molecule inhibitors to the mutated cysteine^10^. Multiple G12C-specific inhibitors are being developed, with some of them showing clinical efficacy^11–13^, and have reinvigorated efforts to find novel ways to inactivate oncogenic KRAS. Mutant-selective inhibitors of KRAS^G12D^, KRAS^G12R^, and other mutants have been recently reported^14,15^. Additionally, pan-RAS inhibitors that target classical RAS isoforms, H-, N-, and KRAS, have also been developed and have entered clinical trials^11,16,17^. Thus, small molecule targeting of RAS is now possible through multiple different chemical approaches. A detailed understanding of possible mechanisms by which oncogenic KRAS can be inactivated may facilitate additional therapeutic efforts to effectively disrupt oncogenic activity.

Deep mutational scanning (DMS) is an approach that utilizes massively parallel sequencing to measure the functional impact of many variants of a protein simultaneously in a single experiment. We have previously utilized this approach to identify resistance mutations to KRAS^G12C^ inhibitors as well as to define structure-function relationships in the SHOC2 protein^18–20^. In addition, recent DMS studies of HRAS have demonstrated the critical context dependence of mutational impact on the functional activity of the RAS protein. Specifically, DMS studies of HRAS in a bacterial two-hybrid selection strategy showed distinct patterns of mutational tolerance for RAS in the presence or absence of a GTPase activating protein (GAP) or guanine nucleotide exchange factor (GEF)^21^. A recent DMS study in yeast also examined the effects of over 26,000 mutations on KRAS folding and effector interactions, defining four major allosteric surface pockets of KRAS^22^. Moreover, DMS experiments for HRAS in the Ba/F3 murine pro-B-cell line identified potential activating mutations (in non-dominant residues) outside of cancer hotspots of RAS by decreasing protein stability and increasing spontaneous nucleotide exchange^21,23^. The impact of mutations in wild-type or oncogenic mutants KRAS expressed in immortalized epithelial or transformed cancer cell contexts has not yet been evaluated.

Here, we utilize DMS to systematically evaluate the impact of nearly all possible missense mutants of oncogenic KRAS in human cell models. We perform DMS on a wild-type (WT) allele of KRAS in a gain-of-function screen and execute a loss-of-function, second-site suppressor DMS screen using a backbone of the oncogenic KRAS^G12D^ allele. We define a model describing the clinical mutation frequency of KRAS as a function of phenotypic selection and cancer cell mutational processes. Moreover, through biochemical and structural analysis of variants that functionally impair oncogenic KRAS, we identified key genetic routes to inactivate the KRAS^G12D^ protein. These studies improve our understanding of clinically relevant *KRAS* variants and provide insights into oncogenic mechanisms that could be targeted in KRAS-mutant cancers.

## Results

### Landscape of clinically observed oncogenic KRAS variants is shaped by transformation potential and mutational processes

We generated a DMS library of WT KRAS-4B (hereafter referred to as KRAS WT library), the predominant isoform of *KRAS*, including a total of 3,536 variants with an average of 18.9 substitutions per position, excluding the first methionine residue. To perform positive selection screening for gain-of-function variants, we employed a transformation model system using the HA1E immortalized human kidney epithelial cell line expressing the SV40 large T- and small T-antigens and the catalytic subunit of telomerase^24^. HA1E cells expressing oncogenic KRAS demonstrate survival on ultra-low attachment tissue culture plates, whereas parental HA1E cells fail to proliferate in the absence of oncogenic KRAS signaling^25–27^ (Methods). We demonstrated that the low attachment growth assay could robustly differentiate HA1E cells transduced with KRAS^WT^ and confirmed expression as well as MAPK activation (Extended Data Fig. 1A&B). Moreover, we transduced 18 distinct KRAS alleles (G12V, A18D, L19F, T20R, Q22K, N26K, D33E, A59G, E62K, E63K, R68S, P110S, C118S, K147N, T158A, R164Q, K176Q, and WT) as well as negative control plasmids into HA1E cells and simultaneously performed low attachment growth assays and xenograft experiments in immunodeficient mice, demonstrating that the growth potential in low attachment culture conditions correlates with tumorigenicity in nude mice (Extended Data Fig. 1C&D).

The KRAS WT DMS library was transduced into HA1E cells and grown in low-attachment conditions for 7 days. Genomic DNA was isolated, and enrichment and depletion scores were calculated as a log2 fold change (LFC) based on the average allele representations in the ultra-low attachment condition at day 7 over that of day 0 (Extended Data Fig. 2A, Methods). We set a threshold for gain-of-function (GOF) alleles (LFC > 0.68) as >2 standard deviations (sd) above the mean LFC of all variants. We then mapped the GOF alleles to the key structural domains of KRAS, including: G1 motif/P-loop (residues 10-16, G2 motif/switch-I/β2 (residues 28-38), G3 motif/switch-II (residues 59-75), β4 (residues 77-83), β5-G4 motif (residues 116-119), G5 motif (residues 144-147), α5 (residues 155-159), and CAAX motif (residues 185-188) (Fig. 1A). These conserved G1−G5 sequence motifs within RAS proteins are pivotal for nucleotide binding, nucleotide-induced structural alterations, and GTP hydrolysis. The majority of the 86 GOF transforming alleles were found in previously characterized codons involved in nucleotide binding and hydrolysis: G12, G13, Q61, N116, K117, S145, and A146 (Fig. 1A, Extended Data Table 1). Within the P-loop, all G12 substitutions, excluding proline, and all G13 substitutions, except alanine and serine, were transforming (Fig. 1A&B, Extended Data Fig. 2B), consistent with substitutions at these positions known to impair GAP-assisted hydrolysis of GTP by sterically blocking the "arginine finger" of the GAP in the active site^28–30^. In the switch-II region, twelve Q61 variants were identified as transforming (Fig. 1A, Extended Data Fig. 2C). Q61 is thought to orient the catalytic water molecule to initiate a nucleophilic attack on the GTP γ-phosphate, and mutations at this residue are known to hinder GTP hydrolysis^31,32^. The G4 and G5 motifs reside within the allosteric lobe of KRAS and are important for binding guanine bases and ribose of the nucleotide. Mutations in these regions result in constitutively activated KRAS due to increased nucleotide exchange rate^33,34^. Four variants at N116, eleven variants at K117, as well as five variants each at S145 and A146 were transforming (Fig. 1A, Extended Data Table 1).

**Figure 1.**
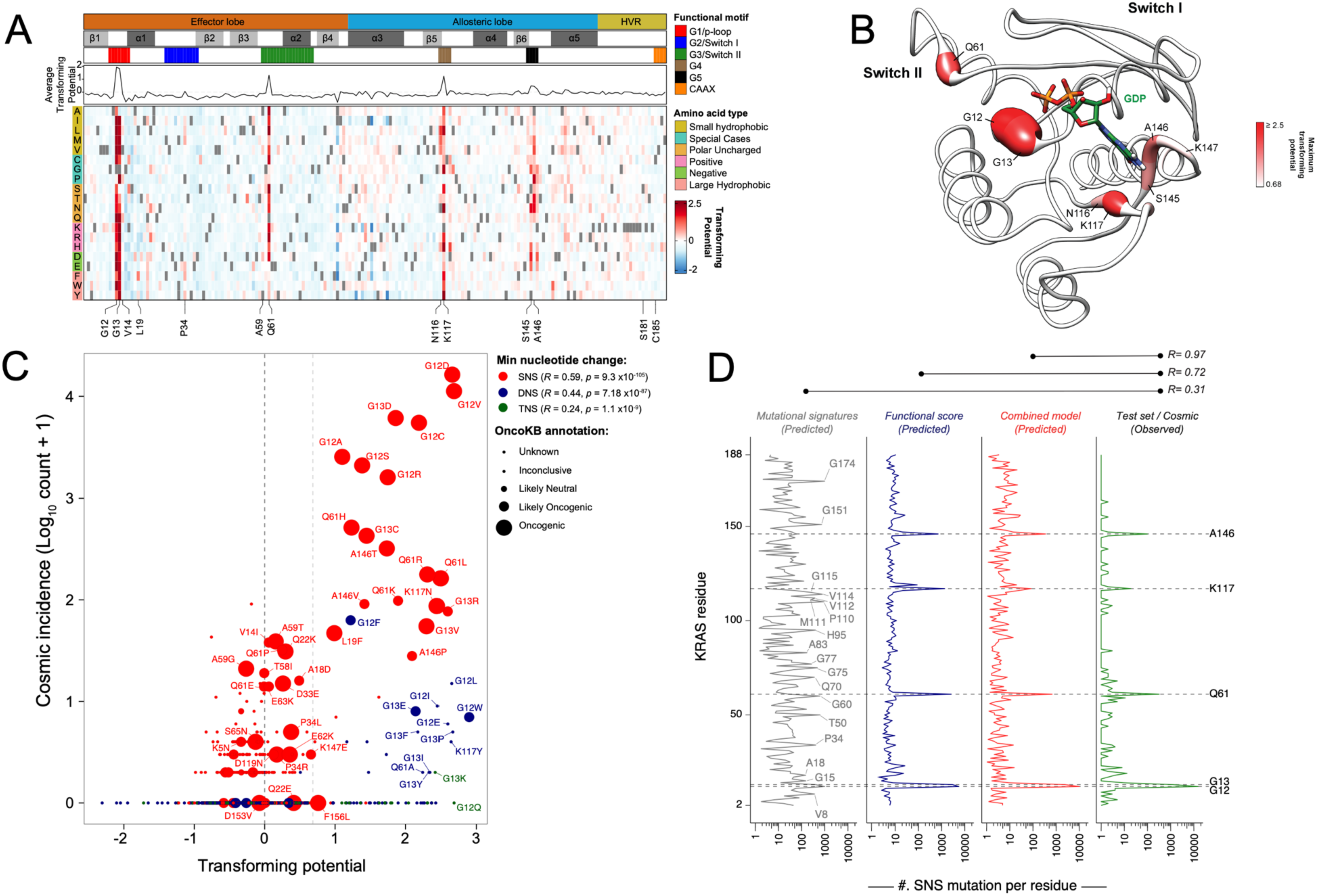
Gain-of-function deep mutational scanning screen highlights KRAS mutational frequency as a function of mutational probability, mutational signatures, and phenotypic selection. (**A**) Heat map representation of LFC allele enrichment (red) and depletion (blue) showing Log_2_ Fold Change (LFC) for each allele from KRAS deep mutational scanning (DMS) gain-of-function screen in HA1E cells, comparing Day 0 and Day 7 data. Each column represents an amino acid in KRAS, and each row represents the substituted residue. Grey squares indicate the missing alleles. The secondary structures, the five nucleotide-binding motifs (G1-G5), and the two Switch motifs are annotated on top, followed by a line graph showing the average LFC of all substitutions at each residue in each screen. (**B**) Mapping of maximal LFC on crystal structure of KRAS per residue position. The color indicated the highest LFC of substitutions at each amino acid and the size correlates with the number of high-ranking putative suppressor mutations at each residue. (**C**) Scatter plot of KRAS variants with functional score from DMS (x-axis) and observed frequency in clinical patient samples (y-axis). Color indicates the minimum number of nucleotide substitutions from native germline codon sequence to mutant variant, with single nucleotide substitution (SNS – red), double nucleotide substitution (DNS – blue), and triple nucleotide substitution (TNS – green). Relative size of bubble indicates OncoKB annotation of oncogenicity. (**D**) Poisson distribution model of KRAS single nucleotide substitution (SNS) spectrum as a function of mutational signature and functional impact is presented. Prediction of SNS counts were carried out using the indicated models trained on KRAS single nucleotide variants occurrences in the GENIE dataset and tested on the KRAS SNS variant occurrence from the COSMIC dataset. The mutation-level Pearson correlation coefficient between predicted and observed counts are presented on top.

Other than the previously well-characterized six codons (G12, G13, Q61, N116, K117, and A146), four positions had 2 transforming variants (K16, L23, D119, and K147) and another six codons had 1 transforming variant per position (Fig. 1A, Extended Data Table 1). We identified novel transforming variants, K16S, K16C, L19F, and L23F within the α1-helix (Extended Data Fig. 2D). K16 interacts with the β- and γ-phosphates of GTP, and mutations at this position likely result in increased nucleotide off-rate. L19 is not directly involved with GTP binding. However, the L19 sidechain faces toward the core of the protein, and mutations likely distort the neighboring nucleotide-binding pocket. Finally, L23 has been previously reported to mediate KRAS-RAF1 interactions at the RAF1 cystine-rich domain (CRD)^35^, and mutations here are likely to impact RAF1 binding. The C-terminal hypervariable region (HVR) (residues 165-189) in KRAS contains the polybasic region (residues 175-180), which is important for KRAS membrane association and localization. Furthermore, KRAS requires farnesylation at C185 within the HVR for proper membrane association and anchoring, and this process is followed by the cleavage of the last three residues from the C-terminal CAAX motif (CVIM). Although mutations within the HVR generally exhibited negative LFC scores, only four variants (S181G, T183A, C185T, I187H) scored two standard deviations below the mean (LFC < -0.98) (Extended Data Table 1).

Previously reported weakly transforming variants A59G, G60E, and Q22K^36,37^, as well as several germline variants of KRAS known to cause congenital diseases such as Noonan Syndrome (e.g., V14I, Q22R, T58F/I, N116S, and D153V)^38–40^, did not meet our stringent statistical threshold for GOF alleles in this screening assay (Extended Data Fig. 2E). Yet, we were able to identify other alleles linked to germline pathogenic variants as transforming (N116H/L/V)^40^, as well as moderately transforming alleles (Q22D and G60S)^41,42^ at greater than one standard deviation above the mean (LFC > 0.27) (Extended Data Fig. 2E). Our assay effectively detected strongly transforming alleles but not weak ones, consistent with most germline variants associated with Noonan Syndrome being weakly activating alleles. Conversely, 92 variants scored two standard deviations below the mean (LFC < -0.98) (Extended Data Fig. 2E), and the largest numbers of variants scoring as depleted were observed at positions A83 on β4-strand and H94 within α3-helix.

Our screen also identified several highly transforming alleles (LFC ≥ 2) that were seldom seen in human cancers, such as Q61A, G13K, G12Q, etc. We hypothesized that this may be due to the lower probability of occurrence as these mutants require more than one nucleotide substitution within the same codon. Thus, we stratified KRAS mutants based on whether they could occur by single nucleotide substitution (SNS), dinucleotide substitution (DNS), and trinucleotide substitution (TNS) and evaluated the correlation of the transforming potential from our DMS screen with the incidence rates in human cancers (COSMIC dataset) (Fig. 1C). In the example of the G12 position, substitution to serine, arginine, cysteine, aspartate, alanine, or valine requires only one nucleotide substitution. These six variants are commonly observed in human tumors, with arginine being the least common of these and is found in 1,571 patients in COSMIC. Mutations to glutamate, tryptophan, phenylalanine, tyrosine, leucine, proline, histidine, isoleucine, threonine, and asparagine require the substitution of at least two nucleotides, and these mutants were observed much less frequently in human cancers (<15 samples each). Mutations to glutamine, lysine, and methionine require at least three nucleotide substitutions, and not a single case was reported in COSMIC.

We observed that certain DNS mutants, such as G12F, appeared more frequently than other transforming DNS mutants in the COSMIC database. We tested the hypothesis that these patients may possess a germline single nucleotide polymorphism at this position that consequently results in the requirement for only a single nucleotide alteration to achieve the DNS change from the canonical sequence, but after thoroughly examining matched germline and somatic mutation datasets from 932 TCGA samples with both germline variant calls and a somatic KRAS mutation, we failed to find germline variations at DNS positions with relatively increased incidence counts (Extended Data Table 2). We further expanded this analysis to 394,656 individuals profiled as part of the UK Biobank and identified 36 unique synonymous nucleotide variants in KRAS, none of which were in the codons encoding G12, G13, or Q61 (data not shown). Overall, we found that SNS displayed a greater correlation between transformation potential and incidence rates (R = 0.59) compared to DNS (R = 0.44) and TNS (R = 0.24) (Fig. 1C).

Context-dependent mutational processes have been shown to impact observed frequencies of mutations in human cancer^43–45^. Indeed, recent studies have demonstrated that mutational processes contribute to KRAS mutations in a tissue-specific manner, likely causing their uneven distribution across cancers^46^. We hypothesized that the functional impact of KRAS mutations, along with mutational signatures, would reflect the clinical distribution of observed mutations. To test this, we modeled the clinically observed mutational spectrum of KRAS as a function of mutational signatures observed in lung adenocarcinoma, colorectal adenocarcinoma, and pancreatic ductal adenocarcinoma along with the transforming potential from our DMS screen. Poisson distribution models were trained on somatic *KRAS* mutations in the COSMIC v97 database and validated using the GENIE database^44,47–49^. We found that mutational counts predicted by our DMS screen alone revealed a strong correlation with mutations observed at each codon position in human cancer (R = 0.73), and the addition of mutational signatures to the model improved the prediction of mutation counts (R = 0.97) (Fig. 1D). Our analyses identified an association between the smoking mutational signature (SBS4) and the mutational process underlying the KRAS^G12C^ and KRAS^G13C^ mutations in lung adenocarcinomas (Extended Data Fig. 3)^47^. We noted that, while the function-based model predicted a higher occurrence rate of mutations at Q61, the infrequency of mutational mechanisms linked to Q61 GOF variants drives down the incidence rates (Fig. 1D, Extended Data Fig. 3). Taken together, our findings systematically explain the prevalence of *KRAS* hotspot mutations in human tumors, as a consequence of both functional impact and underlying mutational processes in cancer cells.

### Comprehensive mapping of second-site suppressor mutations that inactivate oncogenic KRAS^G12D^

G12D is the most frequently observed oncogenic KRAS variant in human cancers and has been linked to poorer survival outcomes^8^. To systematically investigate potential mechanisms of inactivation for oncogenic KRAS^G12D^, we conducted a positive selection DMS screen for loss-of-function (LOF) single amino acid mutations in the KRAS^G12D^ oncoprotein. The screen was performed in the HCC827 cell line, an EGFR exon 19 deleted lung adenocarcinoma cell line that has been shown to undergo apoptosis upon hyperactivating MAPK signaling beyond baseline levels^50^. To calibrate the screen results, we used KRAS^G12D/C185D^ as a known inactivating control. Mutations at residue C185 disrupt KRAS farnesylation, preventing its anchoring to the plasma membrane and resulting in inactive KRAS^51^. As expected, expression of KRAS^G12D/C185D^ in HA1E cells was unable to either activate the MAPK pathway or support anchorage-independent growth in the low attachment (Extended Data Fig. 1A & 1B). When LacZ, KRAS^WT^, KRAS^G12D^, or KRAS^G12D/C185D^ was introduced into HCC827, only KRAS^G12D^ induced apoptosis and reduced the population doubling rate (Extended Data Fig. 4A-D). In contrast, suppressor mutant KRAS^G12D/C185D^ did not affect the viability or proliferation (Extended Data Fig. 4D).

To identify second-site suppressor mutations that inactivate oncogenic KRAS^G12D^, we generated a DMS library with a backbone G12D-mutant allele of KRAS-4B (hereafter referred to as KRAS^G12D^ DMS screen, including a total of 3,535 variants with an average of 18.9 substitutions per position, excluding the first methionine residue. We then stably transduced this KRAS^G12D^ DMS library into HCC827 cells, followed by genomic DNA harvesting and sequencing to assess allelic enrichment and depletion over time (Extended Data Fig. 4E). A LOF score was calculated from the relative abundance of sequencing reads of each allele as an LFC of the average of allele representations on day 12 over that of day 0 (Fig. 2A, Extended Data Table 3). In this KRAS^G12D^ LOF HCC827 DMS screen, higher LFC scores corresponded to greater positive selection resulting from more potent inactivation of the oncogenic activity of KRAS^G12D^. We observed multiple putative suppressor mutations that impair KRAS^G12D^ oncogenic activity. As expected, inactivating substitutions at C185 were enriched in the screen (Fig. 2A, Extended Data Table 1&3). While the C185 position proved to be the most mutationally intolerant position, the polybasic region (residues 175-184) was functionally resilient to multiple point mutations (Fig. 2A, Extended Data Table 3). Using C185 as a benchmark threshold for inactivating alleles, we defined the putative suppressor mutations as variants with higher LFC scores in the KRAS^G12D^ HCC827 screen than the weakest inactivating mutation at C185 (LFC > 0.84) (Methods). To achieve an integrative and comprehensive understanding of KRAS LOF variants, we further compared the results from HCC827 suppressor screen to an additional negative-selection KRAS^G12D^ DMS screen performed in HA1E cells (Methods, Extended Data Fig. 5A). Through this analysis, we identified 331 variants that scored in both screens (i.e. intersection of screens) based on C185 benchmark (HA1E LFC < -0.72), with 59 variants exclusively found in the HCC827 screen and 178 mutations that were unique to the HA1E screen (Extended Data Fig. 5B&C). We note that our hit criteria is stringent, as second-site LOF mutations such as G75A and K104Q in the background of KRAS^G12D^ have been previously shown^52^, but score moderately in our screens (Extended Data Table 3). The putative suppressor variants of the HCC827 screen were concentrated around major functional regions within the G-domain (Fig. 2A&B, Extended Fig. 5D). In the HVR, single mutations within the polybasic region were insufficient to inactivate KRAS^G12D^, likely owing to the functional redundancy of the poly-lysine track for membrane association. However, several suppressor mutations were observed within the C-terminal CAAX box. C185 is essential for lipid posttranslational modification and displayed broad intolerance to any substitutions. Additionally, several mutations introducing charged residues at I187 and M188 also caused inactivation (Fig. 2A).

**Figure 2.**
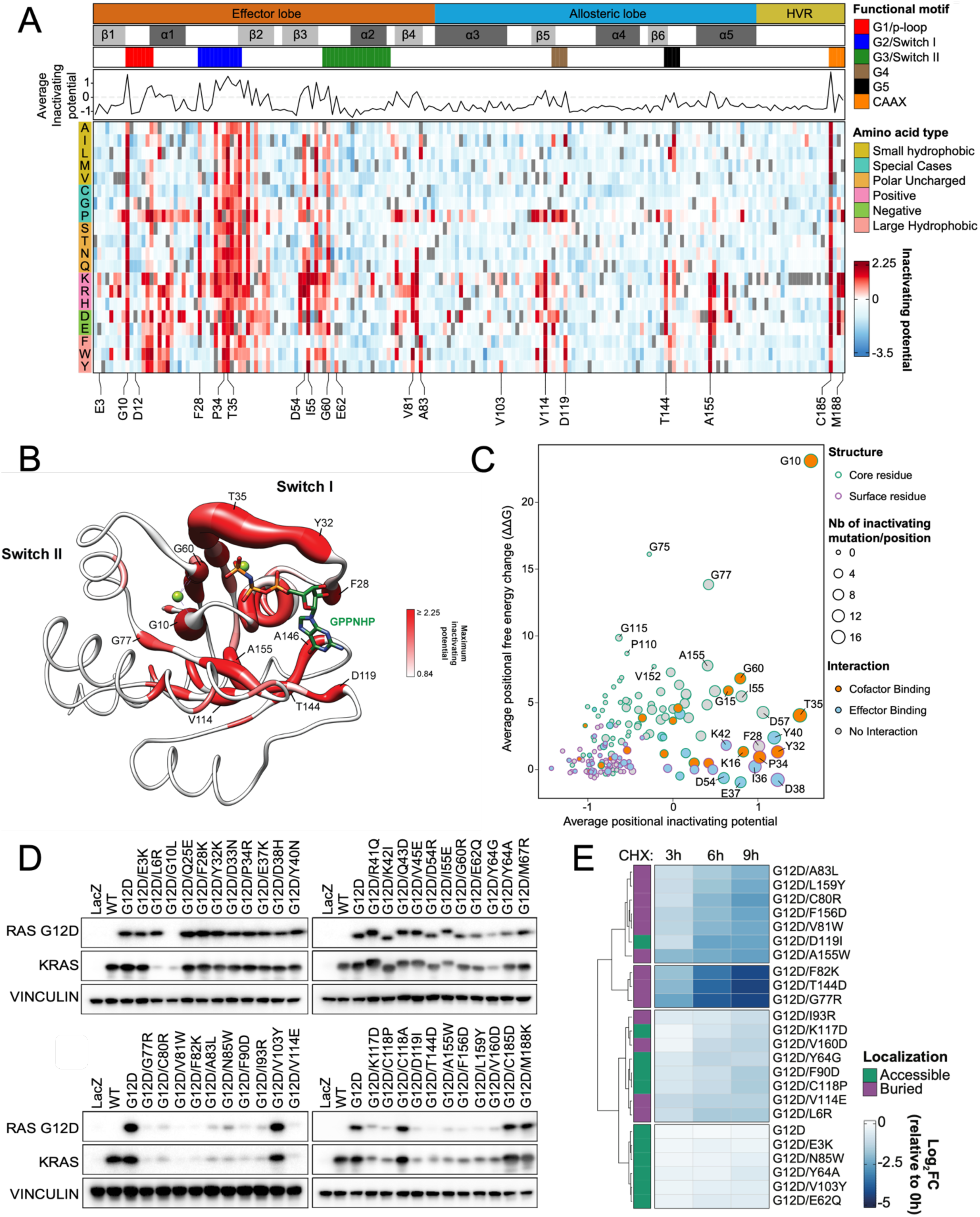
Loss-of-function KRAS^G12D^ screen reveals second-site suppressor mutations and destabilizing mutations. (**A**) Heat map representation of LFC allele enrichment (red) and depletion (blue) showing LFCs for each allele from deep mutational scanning (DMS) screen anchored on KRAS^G12D^ mutant background. The LFC for each variant was calculated based on the Log2 fold change of normalized counts on day 12 compared to Day 0 for HCC827 cells. Each column represents an amino acid in KRAS, and each row represents the substituted residue, and grey squares indicate missing alleles. Secondary structures, the five nucleotide-binding motifs (G1-G5), and two Switch motifs are annotated on top, followed by a line graph showing the average LFC of all substitutions per position. (**B**) Mapping of maximal LFC on the crystal structure of KRAS per residue position. The color indicated the highest LFC of substitutions at each amino acid and the size correlates with the number of high-ranking putative suppressor mutations at each residue. (**C**) Scatter plot showing position-level calculated, mean free-energy change upon mutation (intrinsic KRAS^G12D^ stability) and corresponding average scaled LFC for fitness in the KRAS DMS screen, with higher ΔΔG values corresponding to greater instability and positive DMS LFC indicating inactivating second-site mutation. (**D**) Transient expression of indicated KRAS^G12D^ suppressor mutant alleles in 293T cells. Both RAS^G12D^ and total KRAS were detected. (**E**) Heatmap of Log2FC of RAS^G12D^ levels at indicated timepoints compared to 0 hour following cycloheximide (CHX) treatment in HA1E isogenic cells expressing indicated KRAS^G12D^ mutants.

Out of the inactivating alleles identified in both screens as well as uniquely identified in each screen, we chose 41 alleles covering all secondary structures, motifs, and functional regions for individual validation. We also included Y64A and C118A as two additional LOF controls. Of these variants in the validation set, all but C118A were observed to be functionally inactivating in the HA1E GILA assays and HCC827 growth assays (Extended Data Fig. 5E&F). Furthermore, we selected six alleles and performed HA1E cell line xenograft tumor formation assays, with these experiments showing the expected functional impact on KRAS oncogenicity based on the screening results (Extended Data Fig. 5G). Additionally, we stably transduced these double mutants of KRAS in HA1E and HCC827, and all 41 variants demonstrated reduced levels of phosphorylated(p) MEK and pERK and increased p-STAT3 levels^53^, while the C118A control did not impact downstream signaling (Extended Data Fig. 6&7).

### Identification of destabilizing mutations of KRAS^G12D^ that result in degradation and inactivation

We hypothesized some LOF variants likely induce general protein instability and lower levels of the oncoprotein in cells. To complement our experimental KRAS^G12D^ DMS results, we performed an *in silico* FoldX mutational analysis^54,55^ to determine the predicted mean free-energy change (ΔΔG) upon mutation for each position of KRAS^G12D^ based on a previously reported structure (Methods) (Fig. 2C, y-axis). As anticipated, several solvent inaccessible positions of KRAS^G12D^ are functionally intolerant to mutational change due to predicted destabilization of the protein, with position G10 buried within the P-loop anticipated to exhibit the highest average free energy change upon mutation. Comparing the *in silico* analysis (Fig. 2C, y-axis) with the experimental KRAS^G12D^ DMS screen (Fig. 2C, x-axis), we found that the KRAS positions with high average LFC scores and minimal computationally predicted structural impact were either involved in magnesium/nucleotide cofactor binding or in effector interactions, functions essential for the maintenance of KRAS^G12D^ oncogenic activity. To experimentally test the stability and expression of the selected validation set of 41 LOF mutants, we evaluated baseline expression levels of these KRAS^G12D^ double mutants transfected in 293T cells and observed significant differences in KRAS protein levels (Fig. 2D). About half of the tested mutations led to a reduction in protein levels, especially variants in β4 (77-83), β5 (111-116)-G4 (117-119) and α5 (152-167) regions (Fig. 2D), likely due to destabilization and degradation mediated by the protein quality control system. Lentiviral transduction of these KRAS double mutants in both HA1E and HCC827 also exhibited similar protein expression levels for these mutants (Extended Data Fig. 6&7).

We next examined whether the reduced expression observed on immunoblot for 24 alleles was due to accelerated protein degradation by performing a cycloheximide (CHX) chase assay. Protein levels of KRAS^G12D^ were assessed after 3, 6, and 9 hours of CHX treatment in HA1E cells expressing individual KRAS^G12D^ with second-site mutations (Fig 2E, Extended Data Fig. 8A). For alleles with unclear results, we extended the treatment up to 48 hours and observed faster degradation than KRAS^G12D^ (Extended Data Fig. 8B). For secondary variants such as G10L that may interfere with G12D targeted antibody binding, we devised bicistronic vector that expressed GFP simultaneously with the mutated, HA-tagged KRAS (Extended Data Fig. 8C&D). Overall, we found that half of the selected secondary mutations suppress KRAS^G12D^ oncogenicity by promoting its degradation. When alleles with decreased protein levels were mapped onto the crystal structure of the G-domain of KRAS^G12D^ bound to non-hydrolyzable GTP analog GppNHp (PDB: 6GOF), residues in α3 and α5 helices were orientated toward the core of KRAS. This suggests that substitution at these positions may compromise protein stability, either directly or by hindering helix formation, as mutations of some of these residues to those with lower helical propensity might prevent helix formation and decrease protein stability (Extended Data Fig. 8E).

### Disruption of switch-I/II conformation and effector binding inactivate the oncogenic function of KRAS^G12D^

To further examine mechanisms of inactivation for mutants with preserved protein expression, we selected 14 KRAS^G12D^ second-site mutants (E3K, Q25E, F28K, P34R, R41Q, K42I, Q43D, V45E, D54R, I55E, G60R, E62Q, M67R, and V103Y) from our validation set that represent a broad range of features within the G-domain of KRAS to pursue extensive biochemical and structural studies. We crystallized and solved structures of 11 of the 14 KRAS^G12D^ second-site mutants in complex with GDP/Mg^2+^ (Extended Data Table 4-5). Despite extensive efforts, three KRAS^G12D^ second-site mutants, Q25E, K42I, and Q43D did not crystallize (Extended Data Table 4-5). We proceeded with the structural analysis of these 11 KRAS^G12D^ second-site mutants and performed biochemical analysis on all 14 KRAS^G12D^ second-site mutants to understand the mechanism of inactivation of oncogenic activity (Extended Data Fig. 9A).

We initially performed molecular dynamic (MD) simulation studies on the crystal structures that we resolved and found that average Cα conformational dynamics (RMSD) correlated with the degree of functional loss in the DMS screen (Extended Data Fig. 9B, Extended Data Table 6). In particular, I55E, F28K, and D54R mutants exhibited LFC >2 in the DMS screen and high dynamic motion (overall Cα RMSF > 2.5Å) within the switch regions across the 100ns time course (Extended Data Fig. 9C), consistent with lower melting temperatures compared to KRAS^G12D^ (Extended Data Fig. 9D). I55E and F28K, located on either side of the switch-I region, exhibited a fully open switch-I region. I55 typically resides within a hydrophobic region adjacent to the core b-sheet of KRAS^G12D^, interacting with nonpolar residues within b1-strand and α1/α5-helix. The I55E substitution, introducing a charged/acidic residue, results in a repulsion from the native hydrophobic region and an energetically favorable electrostatic coordination with the Mg ion within the active site. This results in the detachment of the b2-strand from the central b-sheet and the formation of a new antiparallel b-sheet between a newly formed b-strand, consisting of residues preceding the switch-I region, and b2-strand (Fig. 3A). In the case of F28K, substitution with lysine causes a charge repulsion effect with the neighboring K147 residue and a loss of edge-to-face pi-stacking interaction with the guanine moiety of GDP, resulting in an open conformation of switch-I (Fig. 3B). In contrast, D54 resides on b3-strand between the two switch regions and participates in stabilizing hydrogen bond interactions between b2- and b4-strand (Fig. 3C). The D54R mutation causes a charge swap and repulsion of K5 and R41, resulting in new hydrogen bonds with switch-I (S39) and switch-II (D69) and consequent disruption of switch-II α2 helix. Considering the crucial role of the ordered switch conformations of KRAS for effector binding, we hypothesized that mutations that destabilize switch regions, such as I55E, F28K, and D54R, would hinder their ability to bind to effectors. We measured the binding affinity of these double mutants to RAF1-RBD (Ras-Binding Domain), and the mutations resulted in profoundly impaired binding compared to KRAS^G12D^ (Fig. 3D, Extended Data Table 6).

**Figure 3.**
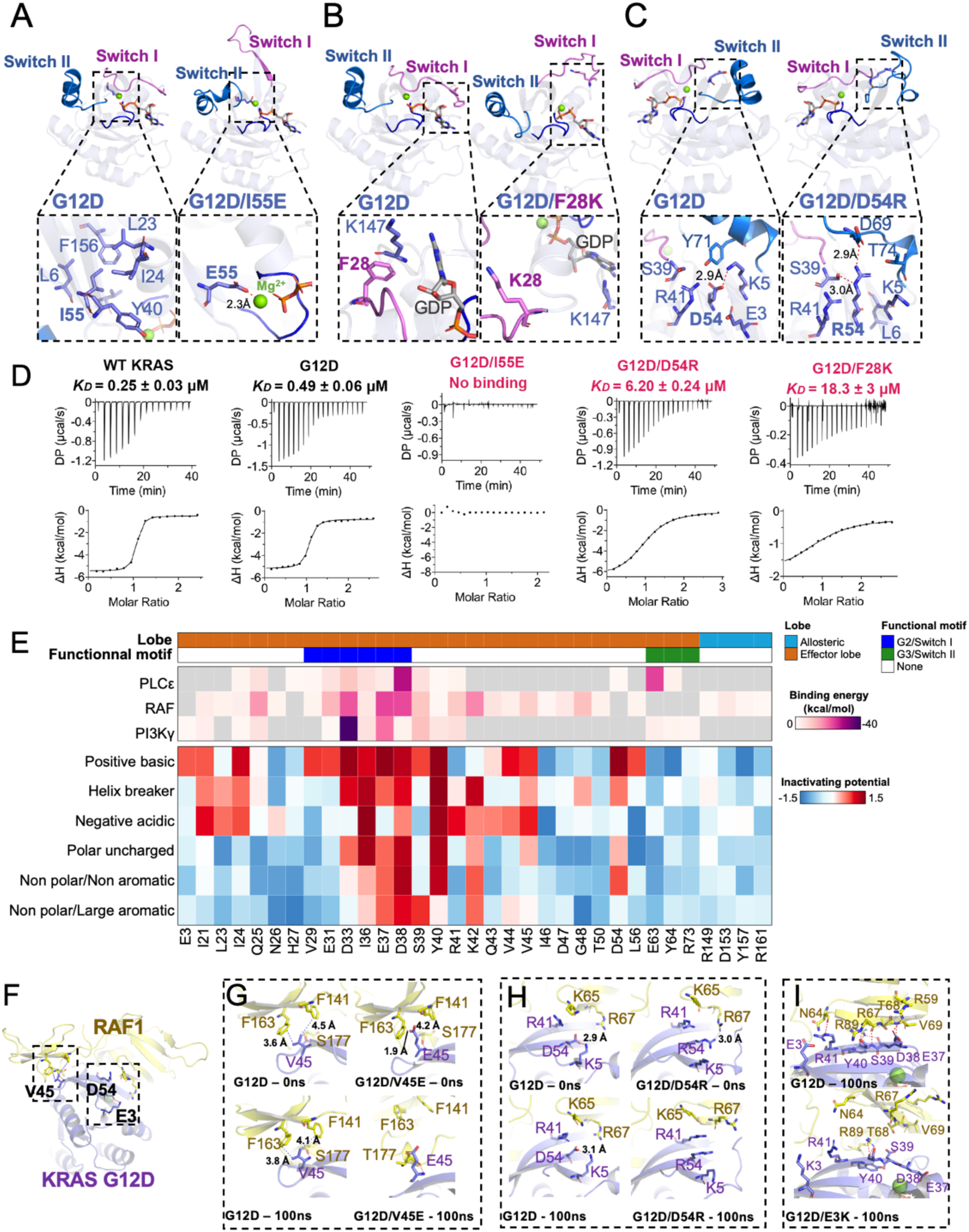
Structural insights and mutational tolerance profiles uncover KRAS^G12D^ inactivation mechanisms by allosteric and orthosteric impacts on switch-I and -II conformations. Structural comparison of GDP-bound KRAS^G12D^ with (**A**) G12D/I55E, (**B**) G12D/F28K, and (**C**) G12D/D54R shows conformational changes in switch-I and -II caused by suppressor mutation. (**D**) Binding affinities (*K_D_* measured by isothermal titration calorimetry) for relevant KRAS^G12D^ inactivating mutants against effector RAF1-RBD, with inactivating mutants labeled in red. (**E**) Heatmap of KRAS effector binding residue interaction energy predicted by Amber10 force-field-based energy calculation (top) and average LFC of residues that have been grouped according to biophysical characteristics, including negative charge (D/E), positive charge (K/R), hydrophobic-aromatic (F/W/Y/H), hydrophobic-small (G/A/V/L/I/M), polar uncharged (S/T/C/Y/N/Q), and helix breaker (P/G). (**F**) Global structural view of KRAS and RAF1(RBD-CRD) with KRAS^G12D^ residues involved in direct RAF1 binding - V45, and proximal residues - E3 and D54 (stick representation) (PDB: 6XHB). Enlarged view of the KRAS-RAF1(RBD-CRD) interaction interface comparing KRAS^G12D^ against (**G**) V45E, (**H**) D54R, and (**I**) E3K.

Next, we assessed the impact of mutations at the effector binding site and proximal residues. We analyzed sidechain substitutions from the DMS screen in terms of their biophysical characteristics and combined this information with binding energies derived from structural data. As anticipated, effector contacting residues within switch-I were generally intolerant to mutational change, in contrast to contacting regions of switch-II, which had minimal functional consequences for most substitutions. Mutations within and proximal to the switch I region (D33, I36, E37, D38, and Y40) resulted in the greatest loss-of-function on average (Fig. 3E), consistent with these residues anticipated to contribute the greatest stabilizing energy for effector binding calculated from previously reported effector bound-RAS structural data by forming interprotein hydrogen bonds and salt-bridges^35,56,57^. We also identified several residues within the interswitch region, with one or more variants scoring as significantly inactivating at positions R41, K42, V44, V45, L53, D54, I55, L56, D57, T58, and A59. For example, K42 demonstrated significant intolerance to any sidechain substitutions except positively charged arginine, which is in line with the KRAS-RAF1 RBD-CRD structure, where K42 forms two hydrogen bonds with CRD residues.

Given that RAS interacts with both RBD and CRD of RAF1, and RAS-CRD interaction is important for full activation of RAF1^35^, we sought to understand a potential mechanism for loss-of-function of these mutants by aligning crystal structures of suppressor mutants to the previously reported RAS-RAF1 RBD-CRD complex structure (PDB: 6XI7) (Fig. 3F). The substitution of V45 with glutamine in KRAS led to decreased affinity for RAF1 RBD-CRD and diminished MAPK signaling, likely due to clashes with residues within the CRD of RAF1^35^. We structurally modeled the impact of mutation at the V45 position by aligning our original V45E crystal structure into a previously reported structure of RAF1 RBD-CRD complexed with KRAS. Following a 100ns MD simulation, we observe a hydrophobic repulsion of KRAS V45E from RAF1 CRD residues F163 and F141 (Fig 3G), consistent with a 2-fold reduction in the binding affinity of KRAS V45E with RAF1 RBD-CRD^10,56–58^. Beyond direct interacting residues, we also sought to structurally understand the impact of proximal mutations on RAF1 binding. D54R resides within the β3 strand and is not directly involved in RAF1 binding. As we previously highlighted, both D54R and E3K mutations result in localized steric hindrance among adjacent side chains within the central β-sheet. In our KRAS-RAF1 RBD-CRD model, we find that D54R causes a direct charge repulsion with R67 of RAF1 (Fig. 3H). Similarly, E3K within the β1 strand also results in a local conformational change due to the E3K shifting the β2 strand, disrupting the KRAS-RAF1 interface (Fig. 3I). These results were further confirmed by measuring binding affinity between KRAS and RAF1-RBD, whereby G12D/D54R, and G12D/E3K resulted in a 12.6-fold and 6.4-fold increase in *K*_D_, respectively, compared to G12D (Extended Data Fig. 10).

The structural superposition of KRAS^G12D^ secondary mutants with HRAS bound to PI3Kγ (PDB: 1HE8) suggests that Q25E, F28K, P34R, and G60R are located near the HRAS-PI3Kγ interface (Extended Data Fig. 9E). Among these secondary mutations, Q25E led to a 2-fold reduction in binding affinity with PI3Kγ, while F28K, P34R, and G60R resulted in complete loss of binding, likely due to mutation-induced conformational changes and loss of critical interactions at the RAS-PI3Kγ interface (Extended Data Fig. 11). Considering that RAS-PI3Kγ interaction involves the switch-II region, G60R mutation abolishes interaction with PI3K but still allows formation of the RAS-RAF1 complex, albeit with reduced affinity. Molecular modelling suggested that G60R in complex with PI3Kγ results in a conformational change in both switch-I and -II due to a new intra-protein hydrogen bond between G60R and E62 (Extended Data Fig. 9F&G). We observe a reduced calculated interaction energy of 19.25 kcal/mol between KRAS^G12D/G60R^ versus KRAS^G12D^, along with an inability to bind PI3Kγ (Extended Data Fig. 9H & 11, Extended Data Table 6). Through rigorous structural and biochemical analysis complemented by molecular dynamics simulations, we have delineated how specific mutations, even those distal from the effector binding site, can mediate structural alterations that significantly impair the interaction of KRAS^G12D^ with key downstream effectors, such as RAF1 and PI3Kγ. Taken together, these studies underscore the intricate interplay between allosteric and orthosteric mutations in dictating the conformational dynamics of switch-I/II regions and subsequent effector binding capabilities of KRAS^G12D^.

### Decreased GEF-mediated nucleotide exchange abrogates KRAS^G12D^ function

The levels of GTP-bound KRAS^G12D^ in cells depend on the ratio of GDP exchange rates mediated by guanine nucleotide exchange factors (GEF), such as Son of Sevenless (SOS), and GTP hydrolysis rates assisted by GTPase-activating proteins (GAP), such as Neurofibromin 1 (NF1). Furthermore, suppression of nucleotide exchange has been shown to abrogate G12-associated oncogenicity^52,58^. To investigate if these suppressor mutations impacted KRAS^G12D^-GTP levels, we measured the intrinsic and SOS-mediated GDP exchange rate and the intrinsic and NF1-mediated GTPase rates.

Among the 14 second-site mutants we selected, 9 exhibited a 3-fold decrease in their SOS-mediated GDP exchange rate (Fig. 4A, Extended Data Fig. 12, Extended Data Table 6). Q25E, P34R, G60R, E62Q, M67R, and V103Y demonstrated a lower SOS-mediated GDP exchange rate than that of the intrinsic GDP exchange rate of KRAS^G12D^. R41Q, Q43D, and D54R were weakly SOS-defective as these had higher exchange rates than the intrinsic rate but lower than the SOS-mediated GDP exchange rate in KRAS^G12D^. Overlaying the KRAS^G12D^ suppressor mutants on the nucleotide-free RAS bound at the catalytic site of the SOS in the RAS-SOS complex showed that all secondary mutations with decreased SOS engagement were located on the RAS-SOS interface. Comparing the structure of RAS-SOS complex with structures of P34R, R41Q, D54R, G60R, E62Q, M67R, and V103Y showed that these mutations resulted in the loss of key interactions with SOS or significant conformational changes rendering them unable to interact with SOS (Fig. 4B&C, Extended Data Fig. 13). Intriguingly, several second-site mutants exhibited a reduction in conformational movement as evaluated by MD simulation. For example, V103 resides on α3 helix at the intersection with switch-II/α2 helix. The substitution of V103 with tyrosine, containing a large aromatic sidechain, sterically blocks the dynamic movement of α2 helix (Extended Data Fig. 14A). This is reminiscent of the effects seen in previous RAS mutants and KRAS inhibitors targeting the switch-II pocket, thereby augmenting the preference for GDP binding^10,59–61^. The P34R secondary mutation likely reduces nucleotide exchange rate due to a new bidentate interaction observed in the crystal structure following MD simulation, stabilizing switch-I toward the α and β phosphates of GDP (Extended Data Fig. 14A). Over a 100ns MD simulation, we observed a structural stabilization of GDP and switch-I movement in V103Y and P34R compared to KRAS^G12D^ mutant structure alone (Extended Data Fig. 14B&C). In contrast, P34R mutation in KRAS^WT^ is GOF (Fig. 1C), and MD simulations of a structural model of KRAS^P34R^reveal a more structurally dynamic and open GDP binding site (Extended Data Fig. 14D), likely increasing the rate of GDP/GTP exchange.

**Figure 4.**
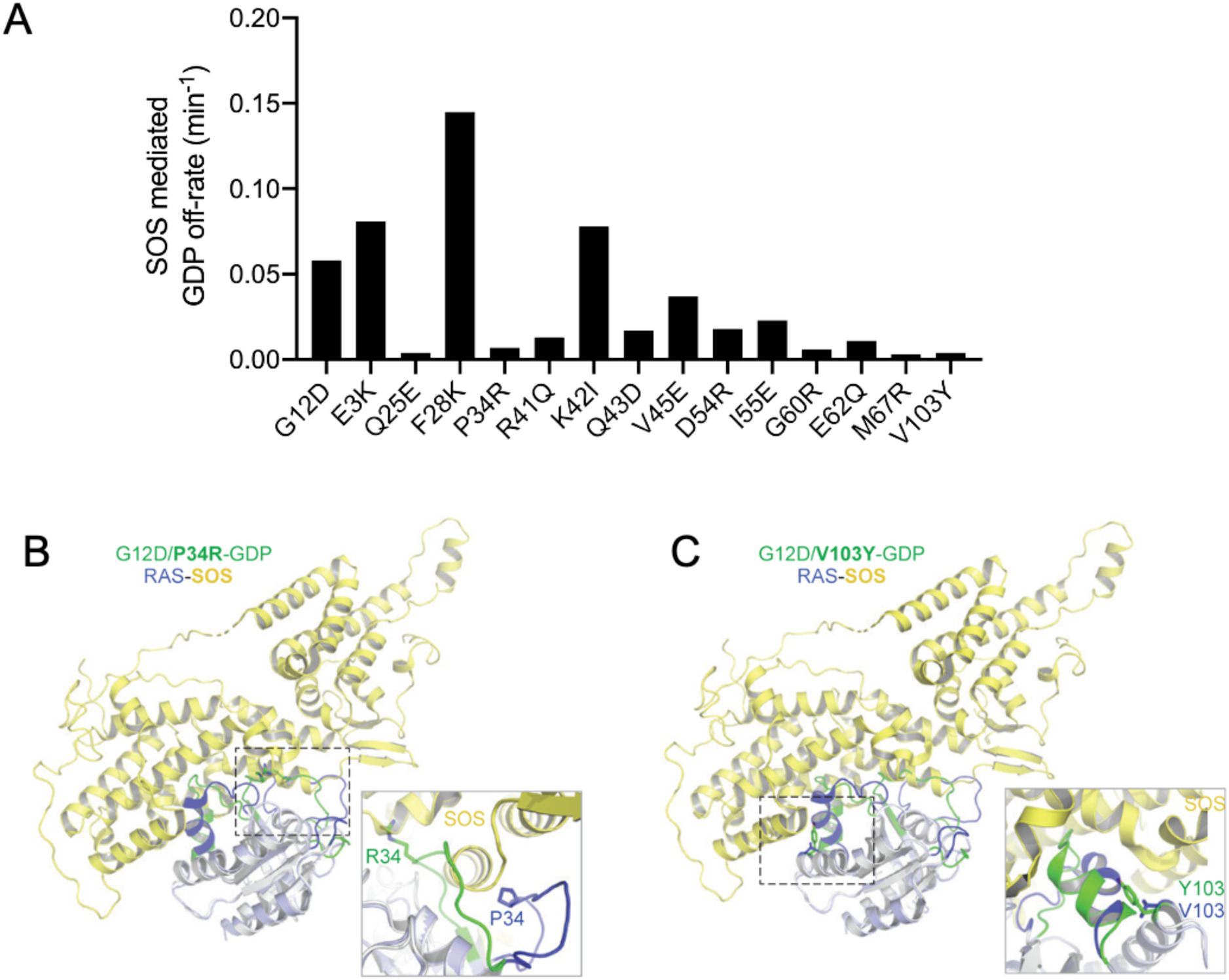
Subset of KRAS^G12D^ inactivating mutations that result in increased GDP engagement through conformational locking and reduced SOS1-dependent GDP exchange. (**A**) SOS-mediated GDP exchange activity: bar graph of SOS-mediated GDP off-rate of KRAS^G12D^ and inactivating mutants. (**B, C**) Superposition of structures of KRAS^G12D^ inactivating mutants P34R (B) and V103Y (C) with HRAS bound at the catalytic site in the HRAS-SOS complex (PDB 1NVW) shows the impact of the inactivating mutation on RAS-SOS interaction. An enlarged view showing the interaction of mutated residues with SOS is shown in the box in each panel. SOS is colored yellow, and regions that undergo significant conformational changes in WT HRAS and KRAS^G12D^ inactivating mutant structures are highlighted in blue and green, respectively. Side-chain atoms of inactivating residue are shown in stick representation.

None of the 14 mutants showed increased intrinsic or NF1-mediated GTPase activity (Extended Data Fig. 15, Extended Data Table 6), suggesting that these suppressor mutations do not enhance GTP hydrolysis in KRAS^G12D^. Binding affinity measurements with NF1-GRD (GAP-related domain) also showed no increased affinity for any of the suppressor mutations (Extended Data Fig. 16, Extended Data Table 6). Notably, the F28K, I55E, and G60R mutants demonstrated a complete loss of both intrinsic GTPase activity and KRAS-NF1 interactions in our assays, indicating a significant impact caused by these secondary mutations. These results suggest that none of the secondary mutations could increase or restore intrinsic or GAP-mediated GTPase activity in KRAS^G12D^. Comparison of the ratio of GDP exchange and GTP hydrolysis rates indicates that Q25E, P34R, R41Q, D54R, E62Q, M67R, and V103Y would have lower GTP levels compared to KRAS^G12D^, and that may contribute to the revertant phenotype.

## Discussion

Through a systematic approach integrating DMS with biological validation and structural analysis, we present a comprehensive structure-function analysis of both gain- and loss-of-function variants of KRAS. We have developed an in-depth model for clinically observed mutational frequencies as a composition of mutational processes and protein function. Furthermore, through systematic genetic studies, we have elucidated the paths to inactivate the KRAS^G12D^ oncoprotein, including impacting protein stability, switch-I/II configuration, effector binding, as well as both intrinsic and SOS-mediated nucleotide exchange activity (Fig. 5). This work provides a framework for interpreting KRAS variants in a clinical setting and a roadmap for exploring therapeutic strategies to inhibit oncogenic KRAS.

**Figure 5.**
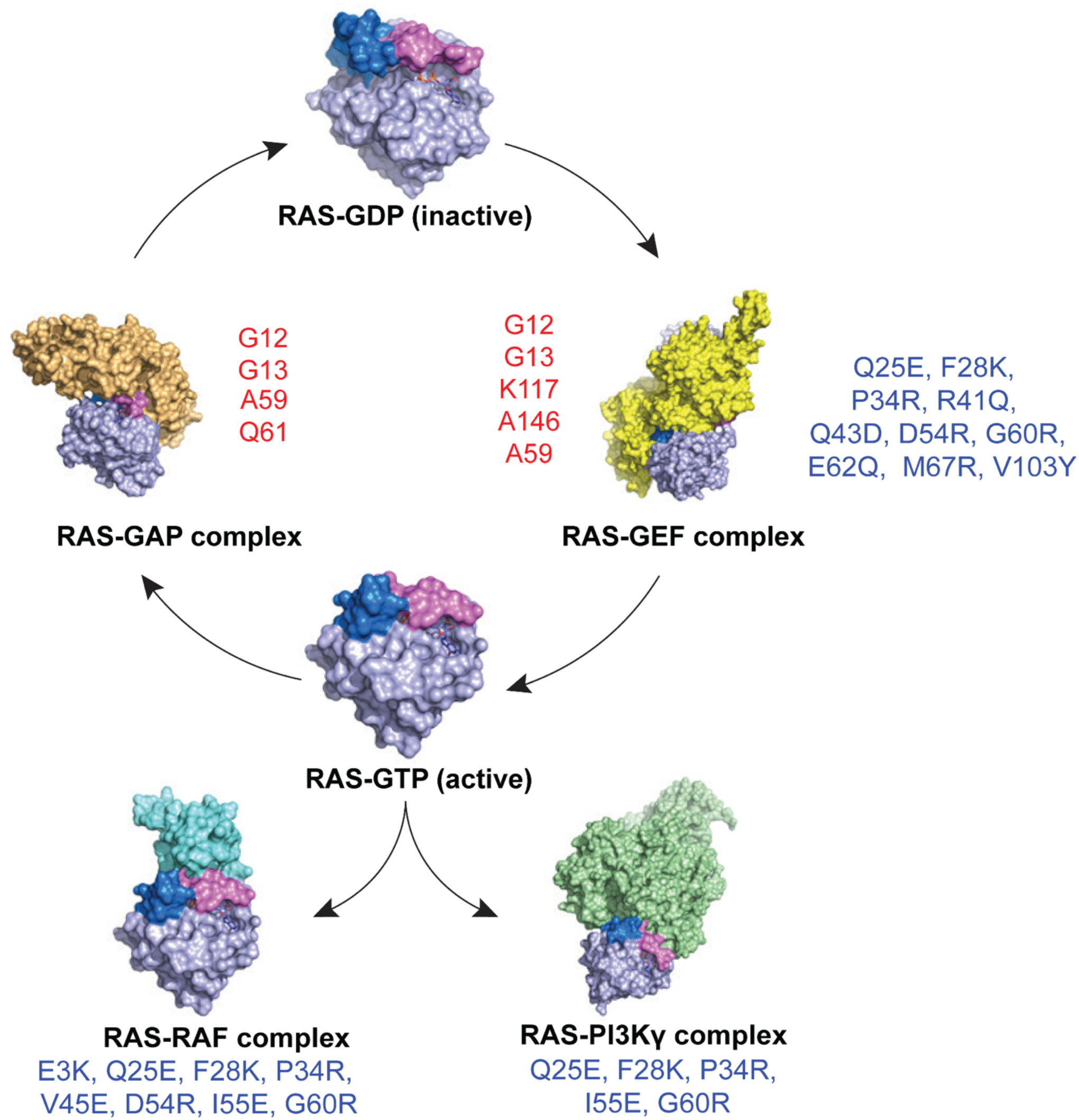
Schematic representation illustrating the impact of second-site inactivating mutations of KRAS^G12D^. Schematic representation of KRAS cycling between its inactive GDP-bound form (RAS-GDP) and active GTP-bound form (RAS-GTP), as well as its interactions with regulatory proteins and effectors. The RAS-GDP state is shown at the top center, transitioning to the RAS-GEF complex (right), which facilitates nucleotide exchange. The GTP-bound RAS engages with downstream effectors, including RAS-RAF and RAS-PI3K. RAS-GAP inactivates RAS by promoting GTP hydrolysis, returning RAS to its GDP-bound state. KRAS gain-of-function mutations (red - G12, G13, A59, K117, A146, and Q61) lead to constitutive activation of RAS. Loss-of-function mutations (blue – E3, Q25, F28, P34, R41, Q43, V45, D54, I55, G60, E62, M67, and V103) disrupt interactions with effectors and regulatory proteins (RasGEFs), resulting in reduced oncogenicity of KRAS^G12D^.

In this study, we offer comprehensive evidence elucidating the varied frequencies of KRAS mutations observed in clinical settings. We also illustrate the relationship between observed RAS mutants and their transformative potential to develop a multivariate model that factors in the probability of acquiring mutations at each codon throughout *KRAS*. The significance of the tissue type and genetic background in determining the functional consequences of various KRAS mutations has been shown to affect clinical outcomes and treatment responses^8,62–69^. We studied the transformation potential of KRAS through DMS using the HA1E immortalized embryonic kidney cell line. The function of a gene depends on its biological context, and a comprehensive understanding of its activity may be better achieved through context-driven, distributional learning^70^. DMS screening in other contexts can be further leveraged to determine how different cell types may affect the transformation capabilities of KRAS variants or to enable a systematic comparison of mutational impacts between RAS family members. Within our gain-of-function screen, we also identified rare tumor variants with high transforming potential at KRAS residues K16, L19, and L23 on the α1 helix, suggesting an important role of these residues in regulating the active RAS levels. Both K16 and L19 reside proximal to the GTP binding pocket, and substitution of these residues may impact nucleotide exchange rate and hydrolysis, like the classic RAS mutations at positions G12, G13, Q61, K117 and A146. However, further investigation is required on the impact of mutations at these sites and their associated mechanisms of activation.

Our KRAS^G12D^ inactivation screen also outlined the molecular pathways leading to KRAS loss-of-function, and intriguingly, several of the variants that we structurally investigated reside within previously identified druggable pockets. Our studies identified mutants such as V103Y near the switch-II cryptic pocket, resulting in the steric locking of switch-II and preferential GDP binding^10,59–61^. Furthermore, E3K and D54R reside near the DCAI pocket^71^. We also found additional mutants, such as P34R under switch-I, that may molecularly cage GDP resulting in KRAS inactivation. Small molecules targeting under switch-I that impact KRAS nucleotide affinity may prove to be an additional therapeutic strategy. Thus, our DMS study, along with recent work by Weng et al.^69^, provides a comprehensive mutational reference map to identify functionally relevant pockets within the KRAS protein. Furthermore, as new cryptic pockets within RAS proteins are identified, our structural analysis will serve as an important resource to understand the functional impact of targeting these areas.

The transforming activity of KRAS^G12D^ was abrogated by mutations that affect SOS-mediated nucleotide exchange. This is consistent with previous observations that KRAS mutants with relatively high intrinsic GTPase rates or slow GDP-off rates, are only partially GTP-loaded. KRAS^G12C^ is the best-known example, but KRAS^G12D^ also appears to require SOS activity for full biological activity. However, wild type RAS proteins are far more dependent on SOS and other GEFs compared with oncogenic RAS mutants. Likely for this reason, drugs that target SOS or SHP2, which is essential for SOS activity, have not been well tolerated in clinical trials, even in the presence of KRAS^G12C^ inhibitors or MEK inhibitors which are known to increase GEF activity.

With several promising KRAS and pan-RAS inhibitors on the horizon and entering clinical trials, the need to effectively classify and understand KRAS mutations has become paramount. Here, we demonstrate KRAS allele frequency is a product of mutational probabilities and functional impact and establish a robust foundation for defining functional, clinically relevant KRAS mutants. The comprehensive functional map of transforming alleles of KRAS presented here will serve as a valuable resource to the clinical oncology community to define oncogenic variants of KRAS that may be targeted by one or more drugs in the increasing array of KRAS inhibitors coming to the clinic. Moreover, as additional mutations in RAS proteins emerge in the context of acquired resistance to mutant-selective KRAS inhibitors, this map will provide a foundation to understand the oncogenic potential of each observed RAS mutation, which may ultimately guide clinical care by influencing the choice of the next lines of RAS-directed therapy. Moreover, given the expansive drug-development efforts ongoing in academia and industry that depend on sound understanding of RAS structure and function, this body of work provides multiple novel datasets and structural models to inform future structure-function studies within the RAS therapeutic community. Lastly, by unveiling the landscape of suppressor mutations of oncogenic KRAS^G12D^, this work also paves the way for the discovery of novel inhibitory mechanisms of KRAS and may inform future therapeutic strategies.

## Methods

### Cell Culture

Cells were tested negative for mycoplasma using the mycoplasma detection kit (Lonza #LT07). The 293T and HCC827 cells were purchased from American Type Culture Collection and cultured in DMEM and RPMI 1640 respectively. HA1E cells were cultured in MEM α (Life Technologies; # 12571071). All media were supplemented with 10% FBS and 1× Antibiotic-Antimycotic (Life Technologies; # 15240062). All cell lines were tested mycoplasma negative.

### Plasmids

pLVET-IRES-GFP and pLVET-HA-K-RasG12V-IRES-GFP were gifts from Aki Manninen (Addgene plasmid # 107139 and # 107140); The latter was used as a template to generate pLVET-HA-K-RasG12D-IRES-GFP (HA-G12D) and pLVET-HA-K-RasG12D/G10L-IRES-GFP (HA-G12D/G10L) with QuikChange Lightning Site-Directed Mutagenesis Kit (Agilent; # 210519). The sequences were confirmed by Sanger sequencing.

### Western blotting

The cells in 2D culture were washed with PBS, followed by the addition of RIPA Lysis and Extraction Buffer (Life Technologies; # 89900) supplemented with 1X Halt Phosphatase Inhibitor Cocktail (Life Technologies; #78420) and 1X Halt Protease Inhibitor Cocktail (Life Technologies; # 87785). The HA1E cell suspension in ultra-low plates were added to tubes with PBS and spun down. The pellets were then resuspended with the lysis buffer. After lysis for 30 minutes on ice, the lysates were centrifuged at 13000 rpm for 15 minutes at 4°C. The supernatant was collected and quantified using a BCA assay (ThermoFisher Scientific # 23225) and protein lysates were then prepared and used for western blotting.

### Antibodies

The antibodies used from Cell Signaling Technology were phospho-p44/42 MAPK (Thr202/Tyr204) (#4370S), p44/42 MAPK (#4695), phospho-Akt (Ser473) (#4060), Akt (#9272), phospho-S6 (Ser240/244) (#2215), S6 (5G10) (#2217), phospho-Stat3 (Tyr705) (#9145S), Stat3 (#4904S), Ras G12D (#14429S) and Vinculin (#4650S). Anti-KRAS rabbit polyclonal antibody was from Proteintech (# 12063-1-AP).

### KRAS WT DMS Screen

Library construction: Lentiviral vector pMT_BRD023 was developed by the Broad Institute Genetic Perturbation Platform (GPP), and library generation was performed similar to as previously described^72^. In brief, A PAC gene is driven by SV40 promotor to confer puromycin resistance. Open Reading Frames (ORF) can be cloned in through restriction/ligation. The ORF expression is driven by EF1a promoter. Cloning: Human KRAS-4B ORF (NCBI Reference Sequence NP_004976.2) was synthesized by GenScript and cloned into pUC57. Missense mutations were created by silicon-based platform developed by Twist Bioscience at a saturation scale. Mutant KRAS-4B cDNA were cloned into pMT_BRD023 with restriction and ligation with NheI and BamHI. We aimed for 1000 colonies per variant, or 4 million colonies for KRAS saturation mutagenesis expression library. Plasmid DNA (pDNA) was extracted from the harvested colonies using Qiagen Maxi Prep Kits. The resulting pDNA library was sequenced via Illumina Nextera XT platform. Lentivirus production: Lentivirus was produced by the GPP at the Broad Institute (Online protocol: http://www.broadinstitute.org/rnai/public/resources/protocols/). Briefly, viral packaging 293T cells were transfected with pDNA library, a packaging plasmid containing gag, pol and rev genes (e.g. psPAX2, Addgene), and VSV-G expressing envelop plasmid (e.g. pMD2.G, Addgene), using TransIT-LT1 transfection reagent (Mirus Bio). Media was changed 6-8 hours post-transfection. Virus was harvested 30 hours post-transfection.

### Screens using the Growth in Low Attachment (GILA) in HA1E cells

Cells were transduced with the pooled lentiviral library with a final concentration of 8µg/mL polybrene and a virus volume to achieve 30% infection efficiency. Next, cells were selected with 1µg/mL puromycin for five days and allowed to recover for two days. On Day 0, recovered cells were either snap frozen for early time point or seeded at 1.0×10^6^ cell density in an ultra-low attachment 10 cm plate (Corning #3262). Cells were spun down and snap frozen on Day 7 and Day 14. For the Day 14 timepoint, the culture medium was refreshed on Day 7. Cell pellets were stored at -80° C until genomic DNA was extracted.

### KRAS^G12D^ DMS screen

Library construction: The vector used for expression of KRAS alleles was the lentiviral vector pMT_BRD023 previously described. Cloning: The KRAS4B reference protein sequence (NP_004976.2) was used as a template to design codon-optimized cDNA sequence. Residue at position 12 was changed from G (Glycine, codon: GGT) into D (Aspartate, codon: GAT). The final ORF was KRASG12D (GGT>GAT) with 6 silent mutations (between 330-348) which were designed for allele-specific knockdown experiments. The sequence of the ORF is as shown below while the G12D mutation and synonymous mutations were underlined.

ATGACTGAATATAAACTTGTGGTAGTTGGAGCTGATGGCGTAGGCAAGAGTGCCTTGACGATACAGCTAATTCAGAATCATTTTGTGGACGAATATGATCCAACAATAGAGGATTCCTACAGGAAGCAAGTAGTAATTGATGGAGAAACCTGTCTCTTGGATATTCTCGACACAGCAGGTCAAGAGGAGTACAGTGCAATGAGGGACCAGTACATGAGGACTGGGGAGGGCTTTCTTTGTGTATTTGCCATAAATAATACTAAATCATTTGAAGATATTCACCATTATAGAGAACAAATTAAAAGAGTTAAGGACTCTGAAGATGTACCCATGGTGCTGGTCGGCAACAAATGTGATTTGCCTTCTAGAACAGTAGACACAAAACAGGCTCAGGACTTAGCAAGAAGTTATGGAATTCCTTTTATTGAAACATCAGCAAAGACAAGACAGGGTGTTGATGATGCCTTCTATACATTAGTTCGAGAAATTCGAAAACATAAAGAAAAGATGAGCAAAGATGGTAAAAAGAAGAAAAAGAAGTCAAAGACAAAGTGTGTAATTATGTAG

This ORF was further flanked at N-terminal with a NheI restriction site and Kozak sequence (GGTTCAAAGTTTTTTTCTTCCATTTCAGGTGTCGTGAGGCTAGCGCCACC) and at C-terminal with a BamHI /MluI restriction site (GGATCCCGGGACTAGTACGCGTTAAGTCGACAATC). Missense mutations were created using the KRASG12D ORF described above by Twist Bioscience as linear fragments of the full-length ORF flanked with adapters for cloning. The fragment library was first digested overnight with NheI and BamHI, then underwent ligation with the pMT_BRD023 lentiviral vector pre-processed with the same restriction enzyme pair. The ligation was carried out with a 5:1 insert-to-vector molar ratio, using a T7 DNA ligase at room temperature for 2h. The ligation was cleaned up with isopropanol precipitation and the resulting DNA pellet was used to transform Stbl4 bacterial cells. Plasmid DNA (pDNA) was extracted from the harvested colonies using a QIAGEN Maxi Prep Kit. The resulting pDNA library was sequenced via Illumina Nextera XT platform to determine the distribution of variants within the library.

gDNA extraction and ORF amplification: gDNA was extracted as previously described. The integrated ORF in the gDNA was amplified by PCR. The PCR products were shot-gun sheered with transposon, index labeled, and sequenced with next-generation sequencing technology. The PCR primers were designed in such way that there is a ∼100 bp extra sequence at each end leading up to the mutagenized ORF region. For the pMT_BRD023 vector, we use these 2 primers:

Forward: 5’-ATTCTCCTTGGAATTTGCCCTT-3’

Reverse: 5’-CATAGCGTAAAAGGAGCAACA-3’

PCR reactions were set up in 96-well plates according to the optimized PCR condition and Q5 DNA polymerase (New England Biolabs) was used as the DNA polymerase. All PCR reactions for each gDNA sample were pooled, concentrated with a PCR cleanup kit (QIAGEN), and separated by gel electrophoresis. Bands of the expected size were excised, and DNA was purified first using a QIAquick kit (QIAGEN) then an AMPure XP kit (Beckman Coulter). Nextera reactions and sequencing was performed as described in Screen deconvolution section for KRAS WT DMS Screen.

### Second-site suppressor screening with HCC827 cells

HCC827 cells were transduced with the lentiviral KRAS^G12D^ DMS library with low multiplicity of infection (<0.3) to make sure each cell could only be infected with one virus. The polybrene was added at 8 µg/mL during transduction. After puromycin selection at 1 µg/ml and cell recovery, a proportion of cells were harvested as day 0 samples. The remaining HCC827 cells were seeded into high-attachment flasks and were harvested after 12 days in culture. Cell pellets were stored at -80° C until genomic DNA was extracted.

### KRAS^WT^ and KRAS^G12D^ DMS data analysis

Software: The reads were processed with a second-generation variant calling software called AnalyzeSaturationMutagenesis (ASMv1.0)^72^ (downloadable at: https://github.com/broadinstitute/gatk/releases). In this version of the software, reads are evaluated full-length, that is, variants are called in the context of entire read (or read pair). The programmed variant is called when 2 conditions are met: (1) the detection of the programmed codon changes, and (2) the absence of any additional nucleotide variations throughout the entire read or read pair. The output files from the ASMv1.0 software were parsed to tally the sum of counts for variants defined by changes detected in the reads relative to the reference ORF sequence. The parser is also downloadable at the above Github site Ultimately, a data-frame file, whose columns are screen samples, rows are variants, cells are counts, is produced. All subsequent analyses were based on this data file.

### Screen deconvolution

gDNA was extracted from frozen cell pellet using Qiagen Maxi kit (#) as per the manufacturer’s protocol. The open reading frame was PCR amplified and gel purified. NGS libraries were made and sequenced on a HiSeq2500 (Illumina) at 150 base pair-end. Extraction of ORF from gDNA: To maintain clone representation, we need to process enough gDNA extracted from enough cells from each screen replicate. In this study, up to 12 separate PCR reactions were done for each gDNA sample. Each PCR reaction were conducted in a volume of 100 uL, and with ∼ 2.5 µg gDNA. Herculase II (Agilent Genomics) was used as DNA polymerase. All 12 PCR reactions of each gDNA sample were pooled, concentrated with Qiagen PCR cleanup kit, and then purified by 1% agarose gel. The excised bands were purified first by Qiagen Qiaquick kits, then by AMPure XP kit (Beckman Coulter). Following Illumina Nextera XT protocol, for each sample, we set up 6 Nextera reactions, each with 1 ng of purified ORF DNA. Each reaction was indexed with unique i7/i5 index pairs. After the limited-cycle PCR step, the Nextera reactions were purified with AMPure XP kit (Beckman Coulter). All samples were then pooled and sequenced with Illumina Hiseq2500 platform.

#### DMS data processing

Reads were aligned to KRAS^WT^ and KRAS^G12D^ reference sequences for the HA1E and HCC827 screens results, respectively. The abundance of each variant was calculated by the fraction of reads compared to the total reads of all variants for each replicates independently. Variants that were not designed in the original library were removed and only intended variants were included for subsequent analysis. Variants with missing values (NA) for at least 2 out of the 3 replicates for either of the timepoints used to calculate the functional scores were removed from the analysis. Variant abundances were defined as the og_2_ fold change between late timepoint (day 7 Low attachment for the KRAS^WT^ GOF DMS or day 12 for the KRAS^G12D^ LOF DMS) in comparison to the library representation at day 0 and were computed by applying a moderated t-test as implemented in the R package limma (version 3.54.2).

### Three-dimensional Mapping

Functional data was mapped onto the structure of KRASWT bound to GDP (PDB: 4OBE) for the HA1E KRAS^WT^ DMS results and onto KRASG12D bound to GPPNHP (PDB: 6GOF) for the HCC827 KRAS^G12D^ DMS results, using UCSF Chimera. Mapping and graphical display of functional data from the DMS screens were mapped onto the crystal structures of KRAS isoforms using Chimera UCSF. KRASWT bound to GDP (PDB: 4OBE) was used for the mapping of the HA1E KRASWT screen results and KRASG12D bound to GPPNHP (PDB: 6GOF) for the HCC827 KRASG12D screen results. Phenotypes were mapped using the “define by attribute” and “render by attribute” functions of the software. For each residue in the structure, we indicated the maximal transformation score (HA1E KRASWT screen) or maximal suppressor score (HCC827 KRASG12D screen) per position as an intensity of red. The number of substitutions per position producing the respective phenotype were represented as a width gradient using the “Worms” attribute of the software.

### Public data

All public data were downloaded in July 2022.

COSMIC v97 data was accessed on www.cancer.sanger.ac.uk. Both targeted and whole genome screen data were used. GENIE v11.1, TCGA, MSK-IMPACT, MSK MetTropism data were accessed on www.cbioportal.org. gnomAD v2.1.1 data were accessed at https://gnomad.broadinstitute.org/ and ICGC data were from https://dcc.icgc.org/.

### Assignment of mutation probability based on trinucleotide mutation context

Single nucleotide substitution signatures and probabilities across the 96 types of trinucleotide mutation types were obtained from COSMIC. Mutational signatures enriched in pancreatic adenocarcinoma, lung adenocarcinoma and colorectal adenocarcinoma, as previously defined^69^, were combined to create into an aggregate list representing mutational processed enriched in all three aforementioned malignancies (SBS1, SBS2, SBS3, SBS4, SBS5, SBS6, SBS8, SBS9, SBS10a, SBS10b, SBS13, SBS15, SBS17a, SBS17b, SBS18, SBS20, SBS26, SBS28, SBS30, SBS37, SBS40, SBS44, SBS45, SBS51). Next, each base in the KRAS^WT^ cDNA sequence was systematically substituted with every other possible base in silico, and the resulting sequences were translated according to the standard genetic code. Mutational probabilities for each mutation in KRASWT that can be achieved through a SNS were computed by considering the nucleotide in 5’ and 3’ of the substituted nucleotide, as previously described^70^. Next, we fitted three different Poisson generalized linear models on the KRAS mutational occurrences from the Genie database, as follows:

- Mutational signature: glm(Genie ∼ SBS1 + SBS2 + … + SBS51, family = poisson(link = "log"))
- Functional score: glm(Genie ∼ LFC, family = poisson(link = "log"))
- Full model: glm(Genie ∼ LFC + SBS1 + SBS2 + … + SBS51, family = poisson(link = "log")

The fitted models were used to predict the mutation counts for each KRAS mutation in the COSMIC v97 database and performance metrics, including correlation were defined between observed and predicted counts.

### KRAS saturation mutagenesis screening data

Functional scores for all the DMS screens presented in this study are available at https://www.targetkras.com/

### Xenograft transplant

Female nude mice (NCRNU-F) were ordered from Taconic Biosciences. 2×10^6 of HA1E isogenic cells were mixed with Matrigel and injected subcutaneously into nude mice and tumor growth were monitored. The animal experiments were done with the approval of DFCI Animal Care and Use Committee.

### Apoptosis assay

After lentiviral infection and puromycin selection, HCC827 cells expressing LacZ, KRASWT, KRASG12D and KRASG12D/C185D were stained with FITC-Annexin V and 7-AAD following the instruction of the FITC-Annexin V Apoptosis Detection Kit with 7-AAD (BioLegend; # 640922) and the samples were run on a BD LSRFortessa cell analyzer.

### Protein degradation assay

HA1E isogenic cells were seeded into 6-well plates and treated with cycloheximide (CHX) (Sigma Aldrich; # C1988) at 20 µg/ml for up to 9 or 48 hours. Protein lysates were prepared for western blot as previously mentioned. Protein bands were quantified using Image J. The KRASG12D variants protein expressions were then normalized to the loading control (vinculin) and the relative level of each KRASG12D variant was then calculated for each timepoints compared to the 0h timepoint. KRASG12D variants were clustered using Euclidean distance and Complete-linkage clustering method.

### CellTiter-Glo assays

HCC827 isogenic cells were seeded into 96-well plate (Falcon; #353072) and CellTiter-Glo 2.0 Assay (Promega; # G9242) was performed following the manufacturer’s instructions. HA1E isogenic cells were seeded into ultra-low attachment 96-well plates (Corning; # 3474) and CellTiter-Glo 3D Cell Viability Assay (Promega; # G9683) was performed after 7 days.

### Cloning, expression and purification of recombinant proteins

Gateway Entry and expression clones for KRAS4b double mutation variants were created following the methods outlined previously^73^. Gateway Entry and expression clones for human RAF1(52-131) and SOScat were described previously^73,74^. Expression clones were as described for Gly-KRAS4b(1-169) Addgene #159539, Gly-KRAS4b^G12D^(1-169) Addgene #159541, and NF1(1198-1530) Addgene #159579. A Gateway Entry clone for human PIK3CG(144-1102) V223K was generated by standard cloning methods and incorporated an upstream tobacco etch virus (TEV) protease cleavage site (ENLYFQG) and a downstream His6 purification tag. Gateway baculovirus Destination vector pDest-602 was constructed by modifying pFastBac-Dual (ThermoFisher) to include a polyhedrin-driven N-terminal maltose-binding protein (MBP) tag to enhance solubility, and a p10-driven enhanced green fluorescent protein (eGFP) marker. A sequence-validated PIK3CG Entry clone was sub-cloned into pDest-602, and the final baculovirus expression clone was used to generate bacmid DNA via the Bac-to-Bac system using the manufacturer’s instructions (ThermoFisher). Final bacmid clones were PCR-verified and used to generate baculovirus^75^.

All KRAS4b proteins, SOScat, and NF1(1198-1530) were expressed as outlined in Taylor *et al*.^76^ using the Dynamite media protocol (16°C induction). RAF1(52-131) was expressed as outlined in Taylor *et al*.^76^ using the auto-induction protocol. PIK3CG(144-1102)-His6 V223K was expressed using the baculovirus-insect cell expression system following protocols described previously^75^. All KRAS proteins, RAF1(RBD; 52-131), and NF1(GRD; 1198-1530) were purified as described for KRAS(1-169) in Kopra *et al*.^77^ with 1 mM MgCl_2_ used for all non-KRAS purification buffers. Briefly, the expressed proteins of the form His6-MBP-TEV-target, were purified from clarified lysates by IMAC (immobilized metal-ion affinity chromatography), treated with His6-TEV protease to release the target protein, and the target protein separated from other components of the TEV protease reaction by a second round of IMAC. Proteins were further purified by gel-filtration chromatography. Purification of SOScat was previously described^74^. PIK3CG(144-1102)-His6 V223K was purified essentially as described for KRAS in Kopra *et al*.^77^ with minor changes. Specifically, i) the purification buffers were 20 mM Tris, pH 8.0, 300 mM NaCl, 1 mM TCEP until the size-exclusion chromatography step, which used 20 mM Tris, pH 8.0, 150 mM NaCl, 1 mM TCEP, ii) the lysate was amended with 25 mM imidazole which was also the concentration of imidazole in the equilibration buffer of the column in the initial IMAC, and iii) as the final protein retains a His6 tag, it bound to the column in the second IMAC step and eluted at high imidazole but was still resolved from His6-TEV protease due to the latter’s higher affinity for the IMAC resin. The fractions with pure protein peaks were combined, flash-frozen in liquid nitrogen, and stored at −80 °C.

### Melting temperature (T_m_) measurements

Thermal stability of wild-type KRAS, KRAS^G12D^, and KRAS^G12D^ second-site mutant proteins bound to GDP were determined using a real-time thermal cycler. Briefly, the protein replicates were assembled in 1.5 ml tubes by adding 90 µl of protein (concentration: 1 mg/ml in the buffer containing 20 mM HEPES buffer pH 7.4, 150 mM NaCl, 5 mM MgCl_2_, and 1 mM TCEP) and 10 µl of the 10X protein thermal shift dye (ThermoFisher). Each protein replicate of 20 µl was added to 3 wells in the 96-well reaction plate and sealed with adhesive film. The sealed plate was spun at 2000 rpm for 2 minutes and loaded onto Quant Studio 3 real-time thermal cycler. The StepOne™ software was used to operate the thermal cycler to acquire the relative fluorescence unit data by ramping the temperature from 25°C to 99°C at the rate of 0.05°C/sec. The acquired data were analyzed using the Applied Biosystems Protein Thermal Shift™ Software to obtain the end-point T_m_ derivative values. The T_m_ derivative values for each protein sample replicate were exported to Microsoft Excel to calculate the mean T_m_ and the standard deviation.

### Intrinsic and SOS1-mediated nucleotide dissociation assay

KRAS proteins (∼4 mg) were diluted in 20 mM HEPES (pH 8.0), 150 mM NaCl, 1 mM TCEP, 200 mM ammonium sulfate and 20 mM EDTA, and incubated overnight at 4° C with 5 mM MANT-GDP (Invitrogen). Excess nucleotide was removed using a GE FPLC using a HighPrep 26/10 column into a buffer containing 20 mM HEPES (pH 8.0), 150 mM NaCl, 2 mM MgCl_2_, and 1 mM TCEP. The efficiency of the MANT-GDP exchange was determined using native mass spectrometry as described previously^78^. MANT-GDP loaded KRAS protein (1.5 µM) was prepared in 40 mM Tris-HCl (pH 7.5), 150 mM NaCl, 2 mM MgCl_2_, and 1 mM TCEP and a final volume of 3 mL. Reactions were initiated by the addition of 2.5 µM SOS_cat_ (564-1048) and 1.5 mM GDP, and the change in fluorescence signal was recorded using an excitation wavelength of 355 nm and an emission wavelength of 448 nm every 30 seconds in a Horiba Jobin Yvon Fluorolog spectrofluorometer, at room temperature. Dissociation rates were calculated by fitting the data to a single exponential decay using Prism graph fitting software.

### Intrinsic and NF1-mediated GTP hydrolysis using phosphate sensor assay

GDP-bound KRAS proteins (∼2 mg) were diluted into 20 mM HEPES, pH 7.3, 150 mM NaCl, 1 mM TCEP, 2 mM MgCl_2_, 20 mM EDTA and 200 mM ammonium sulphate, and were incubated for an hour at room temperature with a 100-fold molar excess of GTP. After the addition of 20 mM MgCl_2_, the mixture was incubated for another 30 minutes at room temperature. Excess GTP was removed by desalting over three PD MidiTrap G-25 columns. The efficiency of GTP exchange was determined using HPLC as described previously^79^. GTP hydrolysis was measured using the Phosphate Sensor assay (ThermoFisher). Specifically, 3 µM of KRAS-GTP, 4.5 µM Phosphate Sensor and 100 nM NF1 (GRD; residues1198-1530) were combined in 50 mM Tris, pH 7.6, 2 mM MgCl_2_, 150 mM NaCl and 1 mM DTT in a final volume of 40 µL. Measurements performed in the absence of NF1 were included to calculate the intrinsic GTPase rates. Potassium phosphate standards (2-fold, 3 µM to 47 nM) were prepared to calculate the amount of phosphate released after GTP hydrolysis. The assay was run in a Corning® 3540 Low Volume 384-well Black/Clear Flat Bottom Polystyrene Not Treated Microplate. Plates were run on a Perkin Elmer Envision every 20 seconds for the first hour and then read for another 7.5 hours at 90-second intervals at room temperature. GTPase hydrolysis rates were calculated by performing a linear regression fit of the data using Prism graph fitting software.

### Binding affinity measurement using isothermal titration calorimetry

The binding affinities of GMPPNP-bound KRAS^G12D^ second-site mutants with downstream effectors (RAF1-RBD and PI3Kγ) and RasGAP NF1 (GRD) were measured using isothermal titration calorimetry (ITC). We used PI3Kγ-V223K mutant for this experiment as it has been shown to bind to KRAS with higher affinity, allowing us to monitor the effect of second-site mutants on KRAS-PI3Kγ interaction. The purified KRAS proteins were first exchanged to a non-hydrolysable GTP analog, GMPPNP, using the protocol described earlier^73^. KRAS^G12D^ second-site mutants, RAF1-RBD, NF1 (GRD) proteins were dialyzed in a buffer (filtered and degassed) containing 20 mM HEPES (pH 7.3), 150 mM NaCl, 5 mM MgCl_2_ and 1 mM TCEP. For the KRAS and RAF1-RBD ITC experiments, 65 μM of KRAS and 650 μM of RAF1-RBD were placed in the cell and syringe, respectively. For the KRAS and NF1-GAP ITC experiments, 50 μM of KRAS and 500 μM of NF1 (GRD) were placed in the cell and syringe, respectively. For KRAS and PI3Kγ ITC experiments, KRAS-GMPPNP mutants and PI3Kγ-V223K proteins were dialyzed in a buffer (filtered and degassed) containing 20 mM Tris pH 8.0, 150 mM NaCl, 5 mM MgCl_2_, 1mM TCEP. ITC run was performed with PI3Kγ at the concentration of 40 μM in the cell and KRAS at the concentration of 400 μM in the syringe. ITC experiments were performed in a MicroCal PEAQ-ITC (Malvern) at 25 °C using 19 injections of 0.4 μl initial injection and, subsequently, 2.2 μl injected at 150-s intervals. Data analysis was performed based on a binding model containing “one set of sites” using a non-linear least-squares algorithm incorporated in the MicroCal PEAQ-ITC analysis software (Malvern).

### Crystallization and data collection

A total of 14 KRAS^G12D^ second-site mutants bound to GDP were screened for crystallization using the sparse matrix screens. Protein concentration used for crystallization of KRAS^G12D^ second-site mutant were as follows: E3K – 8.2 mg/ml; Q25E – 8.5 mg/ml; F28K – 9.9 mg/ml; P34R – 7.6 mg/ml; R41Q – 11.7 mg/ml; K42I – 9.3 mg/ml; Q43D – 8 mg/ml; V45E – 22.7 mg/ml; D54R – 14.8 mg/ml; I55E - 11.3 mg/ml; G60R – 21.5 mg/ml; E62Q – 20 mg/ml; M67R - 9.6 mg/ml; V103Y – 9 mg/ml. For the mutant proteins that did not yield crystals, additional crystallization screens were performed at different concentrations as follows: G12D/Q25E (13.6 mg/ml), G12D/F28K (14 mg/ml), G12D/P34R (15.7 mg/ml), G12D/K42I (14 mg/ml), G12D/Q43D (18.6 mg/ml), and G12D/M67R (14 mg/ml). Except for Q25E, K42I, and Q43D, we were able to obtain crystallization hits for 11 KRAS^G12D^ second-site mutants. Initial crystallization hits were further optimized by varying pH and precipitant concentration as well as by detergent and additive screens. Diffraction-quality crystals were harvested with 15% PEG (polyethylene glycol) 3350 or 25% glycerol as cryo-protectant in the crystal screen solution. Diffraction data for these crystals were collected on 24-ID-C/E beamlines at the Advanced Photon Source (APS), Argonne National Laboratory. The crystallographic datasets were integrated and scaled using XDS^80^. Crystals of KRAS^G12D^ second-site mutants diffracted to a resolution ranging between 1.22 - 2.51 Å. The crystallization conditions are in Extended Data Table 4, and crystal parameters and the data collection statistics are summarized in Extended Data Table 5.

### Structure determination and analysis

Structures of KRAS^G12D^ second-site mutants bound to GDP were solved by molecular replacement using Phaser as implemented in the Phenix/CCP4 suite of programs^81–83^, with a protein-only version of GDP-bound KRAS^G12D^ structure (PDB: 5US4) as the search model. The initial model obtained from molecular replacement was refined using the program Phenix.refine within the Phenix suite of programs^82^, and the resulting *Fo-Fc* map showed clear electron densities for the GMPPNP nucleotide and KRAS protein. The model was further improved using iterative cycles of manual model building in COOT^84^, automated model building in Phenix.autobuild, and refinement using phenix.refine^82^. Once all amino acids that had interpretable electron density were built, potential sites of solvent molecules were identified by the automatic water-picking algorithm in COOT and phenix.refine. The positions of these automatically picked waters were checked manually during model building. Refinement statistics for the structures are summarized in Extended Data Tables 4 and 5. Figures were generated with PyMOL (Schrödinger, LLC). Crystallographic and structural analysis software support was provided by the SBGrid Consortium^85^.

### *In silico* saturation mutagenesis with FoldX

The *in silico* saturation mutagenesis studies on KRAS that evaluate the protein stability from the perspective of free energy change (ΔΔG) upon mutations were performed using the FoldX^54^. MutateX^86^ was used for automation. The overall process was to systematically mutate each available residue within a protein or a protein complex to all other possible residue types and to predict ΔΔGs utilizing the FoldX energy calculation. The RepairPDB function of FoldX was first applied for energy minimization to modify the protein system to reasonable conformations. The BuildModel function was followed for the computational mutagenesis and reporting ΔΔG values.

### *In silico* modeling and preparation of protein systems

Our crystal structures of the KRAS^G12D^ with secondary mutation E3K (PDB entry: 9C43), F28K (PDB entry: 9C3M), P34R (PDB entry: 9C3N), R41Q (PDB entry: 9C3Q), V45E (PDB entry: 9C3R), D54R (PDB entry: 9C3V), I55E (PDB entry: 9C3L), G60R (PDB entry: 9C3Z), E62Q (PDB entry: 9C41), M67R (PDB entry: 9C3K), and V103Y (PDB entry: 9C40) were prepared before modelling and simulations. The module of Protein Preparation in Schrödinger Maestro^87^ was applied to cap termini, repair residues, optimize H-bond assignments, and run restrained minimizations following default settings. Missing loops were modeled using the Prime Homology Modeling module^88,89^. Specifically, for V45E, loops E37-S39, L56-T58, E76-F78, E98-I100, V103-D108, V125-K128, A130-D132, and S145-K147 were modeled; for D54R, loop E62-D69 was modeled; for G60R, loop Q61-E53 was modeled; and for M67R, loop R59-A65 was modeled. Reported KRAS^G12D^ structures bound to GDP and GMPPNP were acquired from the PDB database with PDB entries of 5US4 and 5USJ^87^. Reported KRAS^G12D^ structures underwent the same preparation procedure. For 5US4, loop G60-Y64 was modeled.

Templates of complex models for SOS-RAS, RAF-RAS, PI3Kg-RAS, and NF1-RAS were acquired from the PDB database with PDB entries of 1NVW^90^, 6XI7^55^, 1HE8^56^, and 6OB2^74^. For the SOS in 1NVW, missing loops N26-A31 and R179-G184 were modeled also using the module of Prime Homology Modeling. For the RAF1 in 6XI7, a missing loop E104-G107 was modeled. To build the model of SOS-KRAS^G12D/P34R^ complex and SOS-KRAS^G12D/V103Y^ complex, in silico mutations of P34R and V103Y were realized using the module of 3D Builder in Schrödinger Maestro. To build the model of SOS-KRAS^G12D^, the chain R of 1NVW was replaced with the KRAS in 5US4. Energy minimization in the 3D Builder was applied to avoid collisions between amino acids after the replacement. The same process was applied to build models of SOS-KRAS^G12D/P34R^ complex and SOS-KRAS^G12D/V103Y^ complex, using our crystal structures of KRAS^G12D/P34R^ and KRAS^G12D/V103Y^ to replace the chain R of 1NVW. Energy minimization was followed to avoid collisions between amino acids after the replacement. The same process was applied to build models of RAF1-KRAS^G12D/V45E^ complex and RAF1-KRAS^G12D/D54R^ complex, using our crystal structures of KRAS V45E and D54R to replace the chain A of 6XI7. To build the model of PI3Kg-KRAS^G12D^, the chain B of 1HE8 was replaced with KRAS^G12D^ in 5USJ. Energy minimization was followed to avoid collisions between amino acids after the replacement. The same process was applied to build models of PI3Kg-KRAS^G12D/G60R^ complex using our crystal structure of KRAS^G12D/G60R^ to replace the chain B of 1HE8. To build the model of NF1-KRAS^G12D^, the chain C of 6OB2 was replaced with the KRAS^G12D^ in 5USJ. Energy minimization was followed to avoid collisions between amino acids after the replacement. The same process was applied to build models of NF1-KRAS^G12D/E3K^ complex and NF1-KRAS^G12D/V45E^ complex. All complex models underwent the same protein preparation procedure as described above.

### MD simulations and data analysis

The Schrödinger Desmond MD engine^91^ was used for simulations as described in our previous applications^92^. An orthorhombic water box was applied to bury prepared protein systems with a minimum distance of 10 Å to the edges of the protein. Water molecules were described using the SPC model. Na+ ions were placed to neutralize the total net charge. All simulations were performed following the OPLS4 force field^93,94^. The ensemble class of NPT was selected with the simulation temperature set to 300K (Nose-Hoover chain) and the pressure set to 1.01325 bar (Martyna-Tbias-Klein). A set of default minimization steps pre-defined in the Desmond protocol was adopted to relax the MD system. The simulation time was set to 200 ns for each protein system. One frame was recorded per 200 ps during the sampling phase.

Post-simulation analysis was performed using a Schrödinger simulation interaction diagram. A Python-based analysis script analyze_trajectory_ppi.py was used to monitor contacting residue pairs during the MD course. A Python-based analysis script trj_essential_dynamics.py was used to perform the principal component analysis on MD trajectories based on protein C-alpha atoms.

Protein-protein interaction energy was calculated using Molecular Operating Environment (MOE) (2019.01; Chemical Computing Group). In MOE, before calculating the interaction energy, the minimization and optimization of the protein system were performed under the Amber10:EHT force field (https://infoscience.epfl.ch/record/121435/files/Amber10i.pdf) to root-mean-square (RMS) gradient of the potential energy falls below 0.1 kcal mol-1 Å-1. Default tether restraints from MOE were applied to the system. The interface energy calculation between contacting residue pairs was processed using the module of Protein Contacts. Six types of contacts were identified: hydrogen bonds (H-bond), metal, ionic, arene, covalent and van der Waals distance interactions (distance). The proximity threshold was set to 4.5 Å. Atoms separated by more than this distance were not considered to be interacting. The energy threshold was set to −0.5 kcal mol-1 for H-, H-pi and ionic bonds.

## Acknowledgments

This work was supported in part by grants from the Innovation Grant Program at Harvard Medical School (EK), Samsung Scholarship (EK), Lustgarten Foundation (AJA), Dana-Farber Cancer Institute Hale Center for Pancreatic Cancer Research (AJA, WCH), the Doris Duke Charitable Foundation (AJA), Pancreatic Cancer Action Network (AJA), National Institutes of Health National Cancer Institute K08 CA218420-01 (AJA), R01 CA276268 (AJA), K99 CA270290 (JJK), P50CA127003 (AJA, WCH), U01 CA224146 (AJA, WCH), and ACS MRSG-18-202-01 (ALH). AOG is the recipient of a CIHR Fellowship Award (Application Number 430950). This project was funded in part with federal funds from the National Cancer Institute, National Institutes of Health Contract 75N91019D00024. This work used NE-CAT beamlines (GM124165), a Pilatus detector (RR029205), an Eiger detector (OD021527) at the APS (DE-AC02-06CH11357). We thank John-Paul Denson, Matt Drew, Peter Frank, Bill, Gillette, Brianna Higgins, Jennifer Mehalko, Simon Messing, Min Hong, Shelley Perkins, Kelly Snead, and Vanessa Wall from the Frederick National Laboratory for their help in cloning, expression, and purification of recombinant proteins. The content of this publication does not necessarily reflect the views or policies of the Department of Health and Human Services, and the mention of trade names, commercial products, or organizations does not imply endorsement by the US Government.

## Author Contributions

J.J.K., J.D. S.L., E.K., Y.B. designed and executed the study. X.Y. and D.E.R. generated DMS library and S.L. executed DMS screens. J.D., S.L., E.K., S.H.L, performed in vitro and in vivo experiments. J.D., K.S.K., S.M., and J.J.K. performed integrative DMS-patient analyses. D.E. provided recombinant proteins. T.H.T., S.D., and D.K.S. resolved the KRAS X-ray crystal structures. S.D., D.R., and T.J.W. performed biophysical experiments. J.J.K. and Y.B. performed integrative structure-function analyses, structural modeling, and molecular dynamic simulations. A.G.S., D.V.N., and D.K.S. contributed insights on the biochemistry, biophysics, and structural interpretations. A.O.G., B.W., A.R., J.G.D., K.M.H., and F.M. provided scientific insights. J.J.K., J.D., S.L., E.K., D.K.S., W.C.H., and A.J.A. wrote the manuscript. S.L. D.K.S., W.C.H., and A.J.A. supervised the execution of this study. All the authors edited and approved the manuscript.

## Additional Information

and permissions information is available at www.nature.com/reprints.

## Materials and Correspondence

Correspondence and requests for materials should be addressed A.J.A., W.C.H., or D.K.S., S.L.

## Supplementary Information

Supplementary Information is available for this paper.

## Data availability

The coordinates and structure factors for KRAS^G12D^ second-site suppressor mutant structures are deposited in the Protein Data Bank and can be accessed using the following accession codes: G12D/E3K (9C43), G12D/F28K (9C3M), G12D/P34R (9C3N), G12D/R41Q (9C3Q), G12D/V45E (9C3R), G12D/D54R (9C3V), G12D/I55E (9C3L), G12D/G60R (9C3Z), G12D/E62Q (9C41), G12D/M67R (9C3K), and G12D/V103Y (9C40). Primary data are provided with this paper.

## Code availability

Molecular Operating Environment (MOE) and Schrödinger software are publicly available for commercial and non-commercial use.

