## Extended Data Figures for "Comprehensive structure-function analysis reveals gain- and loss-of-function mechanisms impacting oncogenic KRAS activity"

### Extended Data Fig. 1

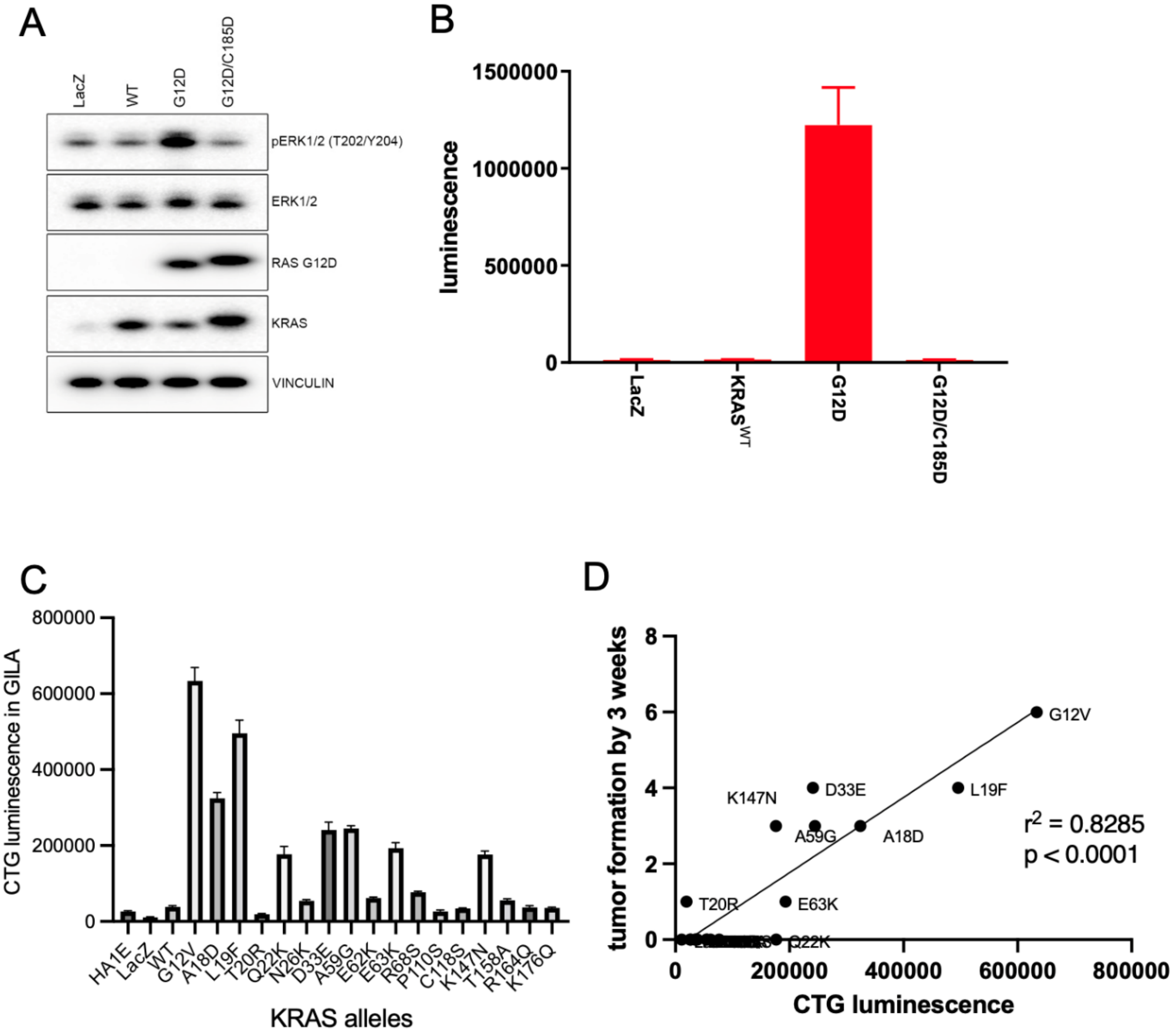


**Impact of KRAS$^{G12D}$ on HA1E growth in ultralow attachment.** HA1E cells were transduced with lentivirus expressing LacZ, KRAS$^{wt}$, KRAS$^{G12D}$ or KRAS$^{G12D/C185D}$ and selected with puromycin at 1ug/ml. (**A**) KRAS expression and downstream signaling were checked. (**B**) 1,000 of HA1E cells stably expressing each allele were grown in ultra-low attachment wells for 7 days followed by CellTiter-Glo 3D assays. Evaluation of HA1E cells stably expressing KRAS WT, G12V, A18D, L19F, T20R, Q22K, N26K, D33E, A59G, E62K, E63K, R68S, P110S, C118S, K147N, T158A, R164Q, and K176Q on (**C**) growth in low attachment and (**D**) *in vivo* growth.

**Extended Data Fig. 2**

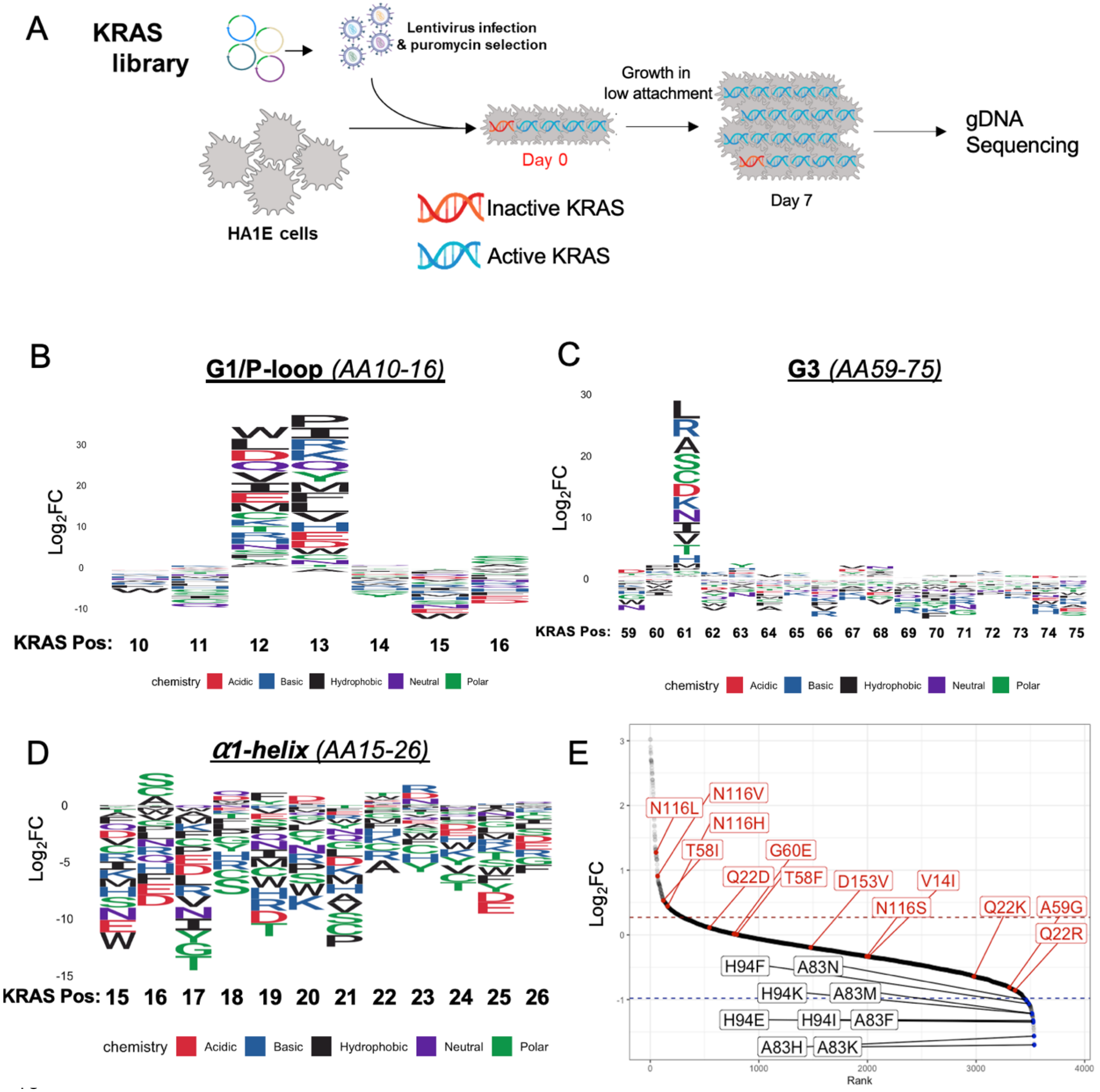


**KRAS Deep Mutational Scanning (DMS) gain-of-function screen.** (A) Schematic overview of KRAS DMS gain-of-function (GOF) screen. Sequence logo plots of KRAS DMS GOF screen results where the height of substituted amino acid indicates the log2

fold change (y-axis) for each amino acid position of KRAS (x-axis) for (B) G1 domain (C) G3 domain, and (D) α1-helix. (E) Waterfall plot of KRAS DMS GOF screen in rank order of each variant functional impact log2 fold change. KRAS variants implicated in Noonan syndrome (red) and select loss-of-function mutants at positions A83 and H94 (black) are indicated.

### Extended Data Fig. 3

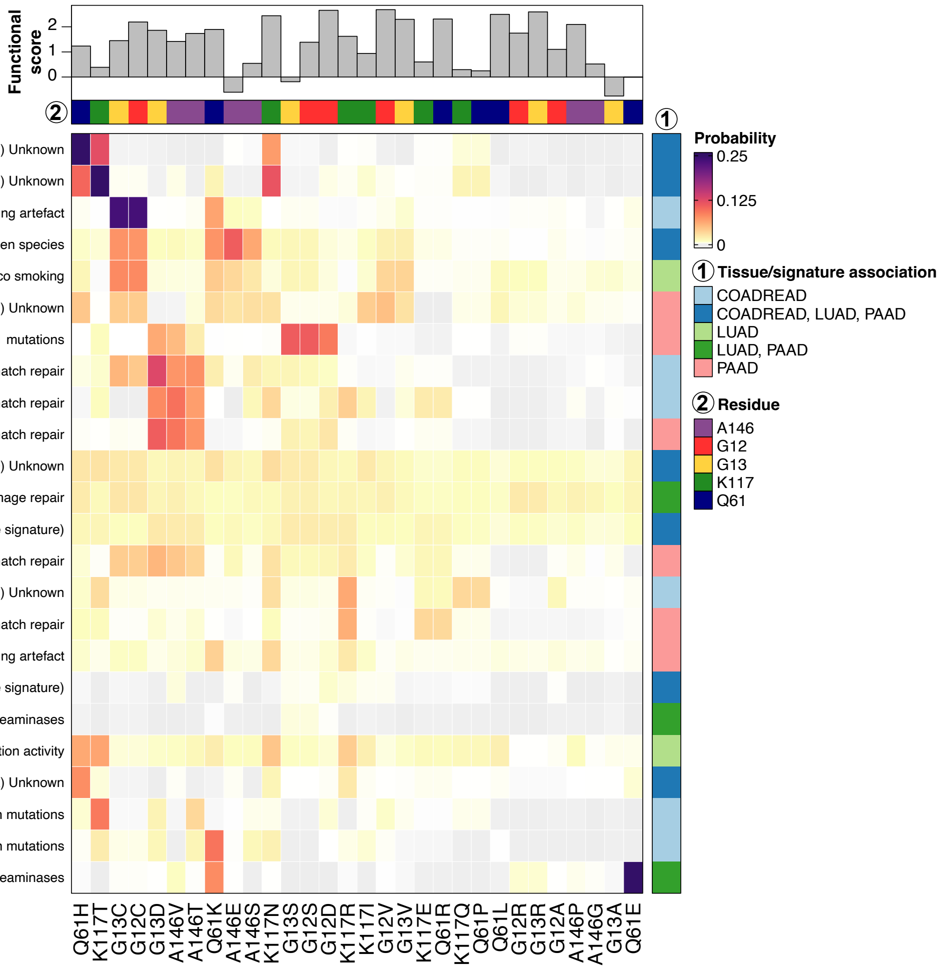


**Oncogenic KRAS variants and their associated mutational signatures.** Heatmap depiction of (1) tissue-specific probability of mutational signatures resulting in (2) a given KRAS mutation. Functional score from HA1E KRAS DMS GOF screen is indicated in barplot above.

**Extended Data Fig. 4**

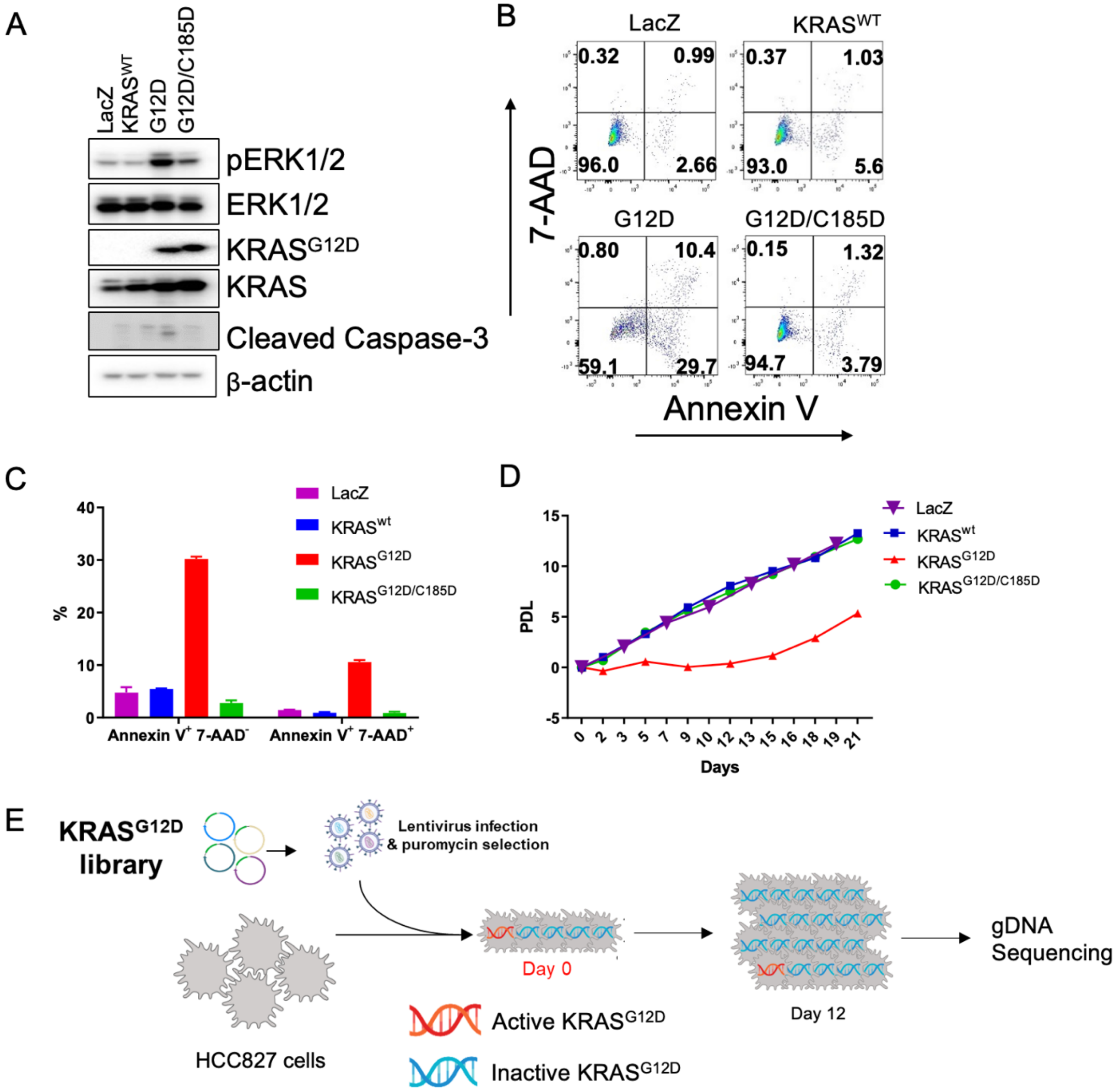


**KRAS$^{G12D}$ Deep Mutational Scanning (DMS) loss-of-function screen.** HCC827 cells were transduced with lentivirus expressing LacZ, KRAS$^{wt}$, KRAS$^{G12D}$ or KRAS$^{G12D/C185D}$. (**A**) KRAS expression and downstream signaling, (**B**) Apoptosis by flow cytometry with FITC-annexin V and 7-AAD dual-labeling, (**C**) Early apoptosis (Annexin V+7-AAD-) and late apoptosis (Annexin V+7-AAD+) are shown. (**D**) Population doubling level (PDL) was monitored for 21 days. One representative of three independent experiments was shown. (**E**) Schematic overview of KRAS$^{G12D}$ DMS loss-of-function (LOF) screen.

61 **Extended Data Fig. 5**

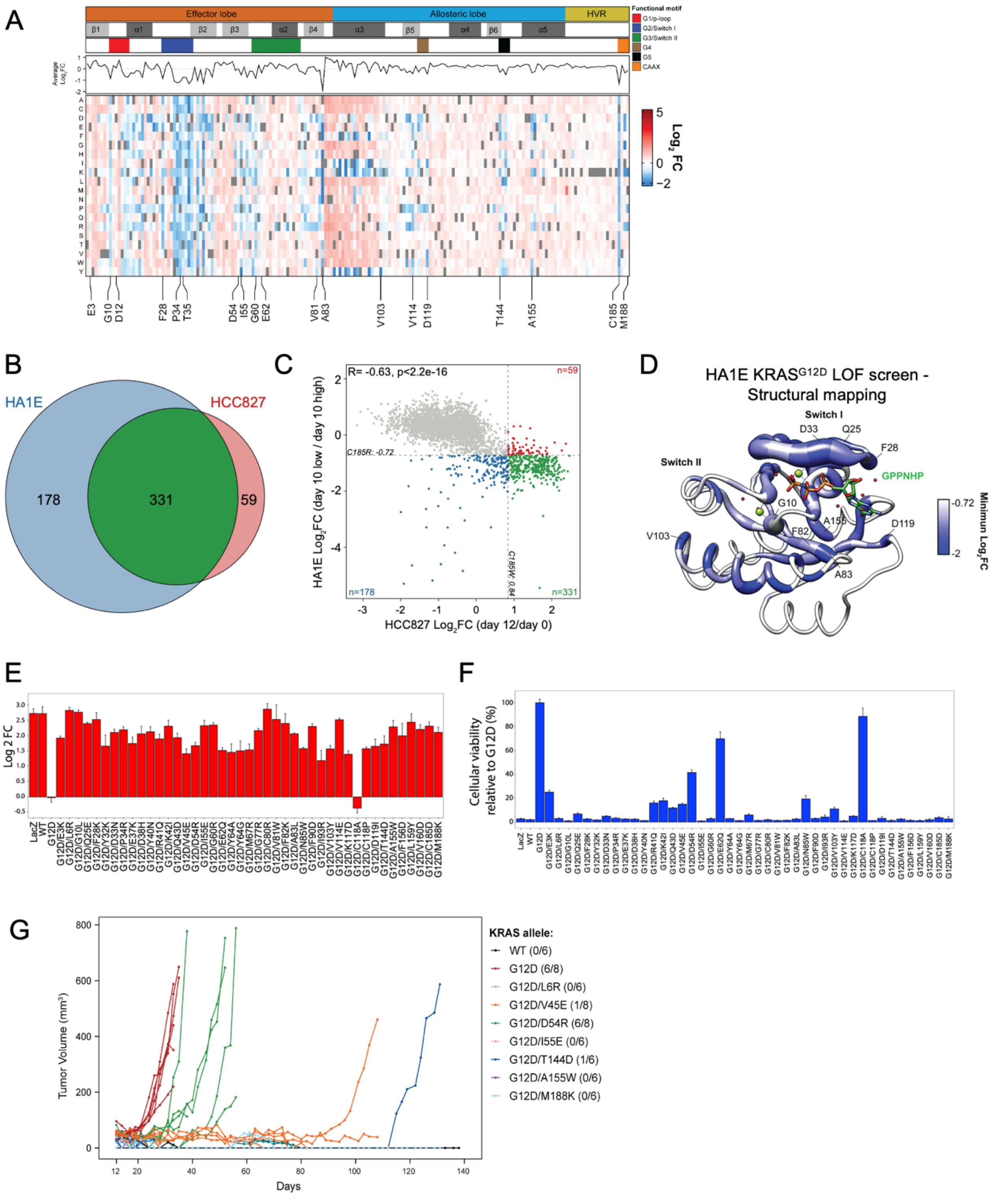


62

**Evaluation of KRAS LOF mutations and validation.** (**A**) Heat map representation of LFC allele enrichment (red) and depletion (blue) showing LFCs for each allele from KRAS$^{G12D}$ deep mutational scanning (DMS) screen. The LFC for each variant was calculated based on the Log2 fold change of normalized counts on the indicated day compared to Day 0 and Day 10 data shown for HA1E cells. Each column represents an amino acid in KRAS, and each row represents the substituted residue, and grey squares indicate missing alleles. Secondary structures, the five nucleotide-binding motifs (G1-G5), and two Switch motifs are annotated on top, followed by a line graph showing the average LFC of all substitutions per position. (**B**) Comparison of putative suppressor mutations against C185 benchmark in KRAS$^{G12D}$ HCC827 screen (red) and KRAS$^{G12D}$ HA1E screen (blue) with a number of overlapping and unique hits indicated in a Venn diagram, defining LOF variants with LFC scores greater. (**C**) Scatterplot of KRAS$^{G12D}$ DMS screen for second site mutants from HCC827 (x-axis) and HA1E (y-axis) screens. C185 benchmark indicated by dashed line, and second site mutant hits that are unique to the HA1E screen (blue), HCC827 screen (red), and overlapping hits (green) are indicated. (**D**) Mapping of maximal loss-of-function observed in HA1E DMS screen on crystal structure of KRAS$^{G12D}$ per residue position. The color indicated the lowest LFC of substitutions at each amino acid and the size correlates with the number of high-ranking putative suppressor mutations at each residue. 43 KRAS$^{G12D}$ suppressor mutants validated in (**E**) HA1E GILA assays, (**F**) HCC827 growth assays, and (**G**) selected variants were evaluated for *in vivo* growth.

**Extended Data Fig. 6**

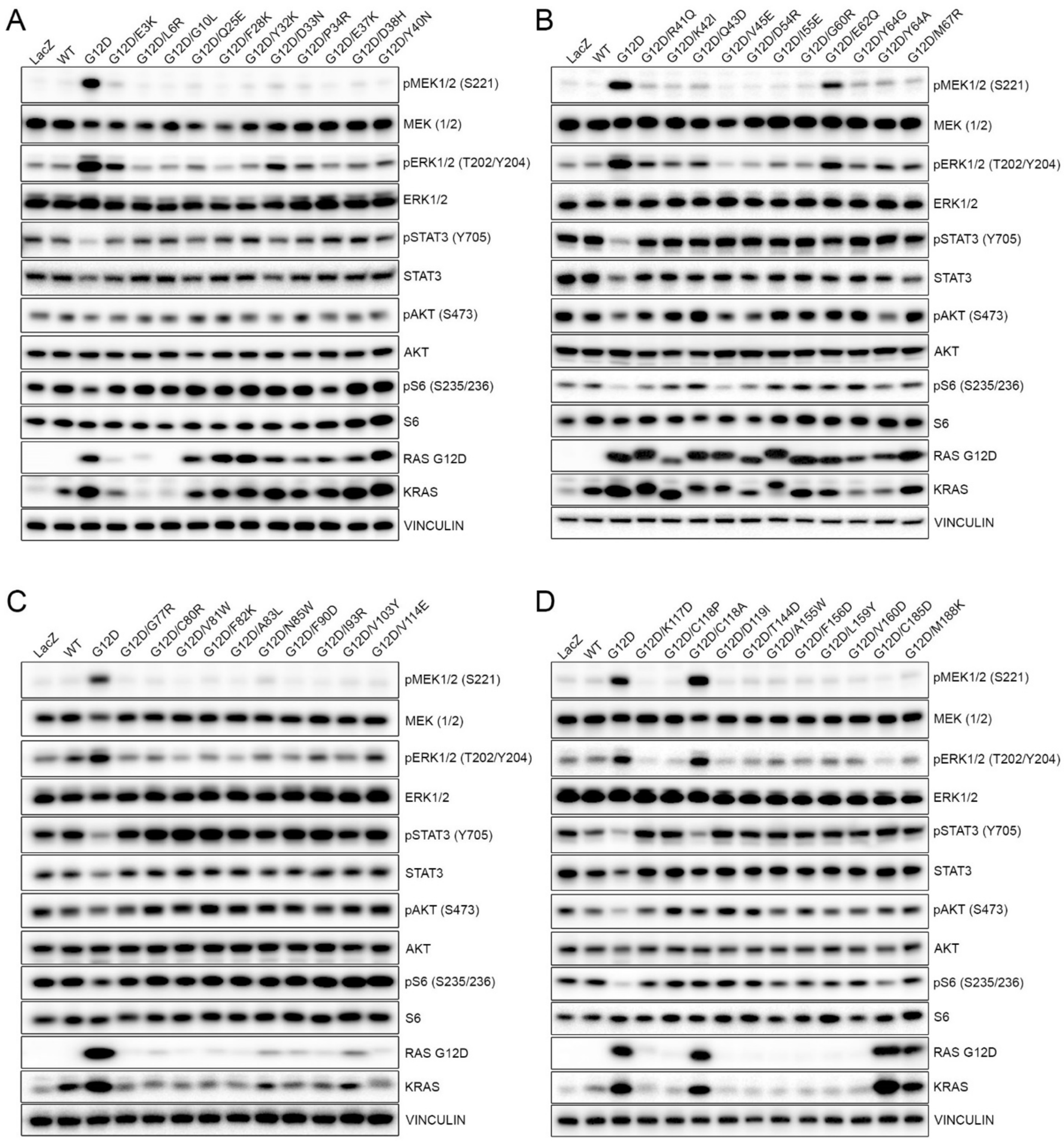


**Signaling pathways in isogenic HCC827 cell lines with KRAS$^{G12D}$ alleles.** HCC827 cells were transduced with lentivirus expressing indicated KRAS$^{G12D}$ alleles. Positive cells were selected with puromycin and main downstream signaling molecules were checked with western blots.

### Extended Data Fig. 7

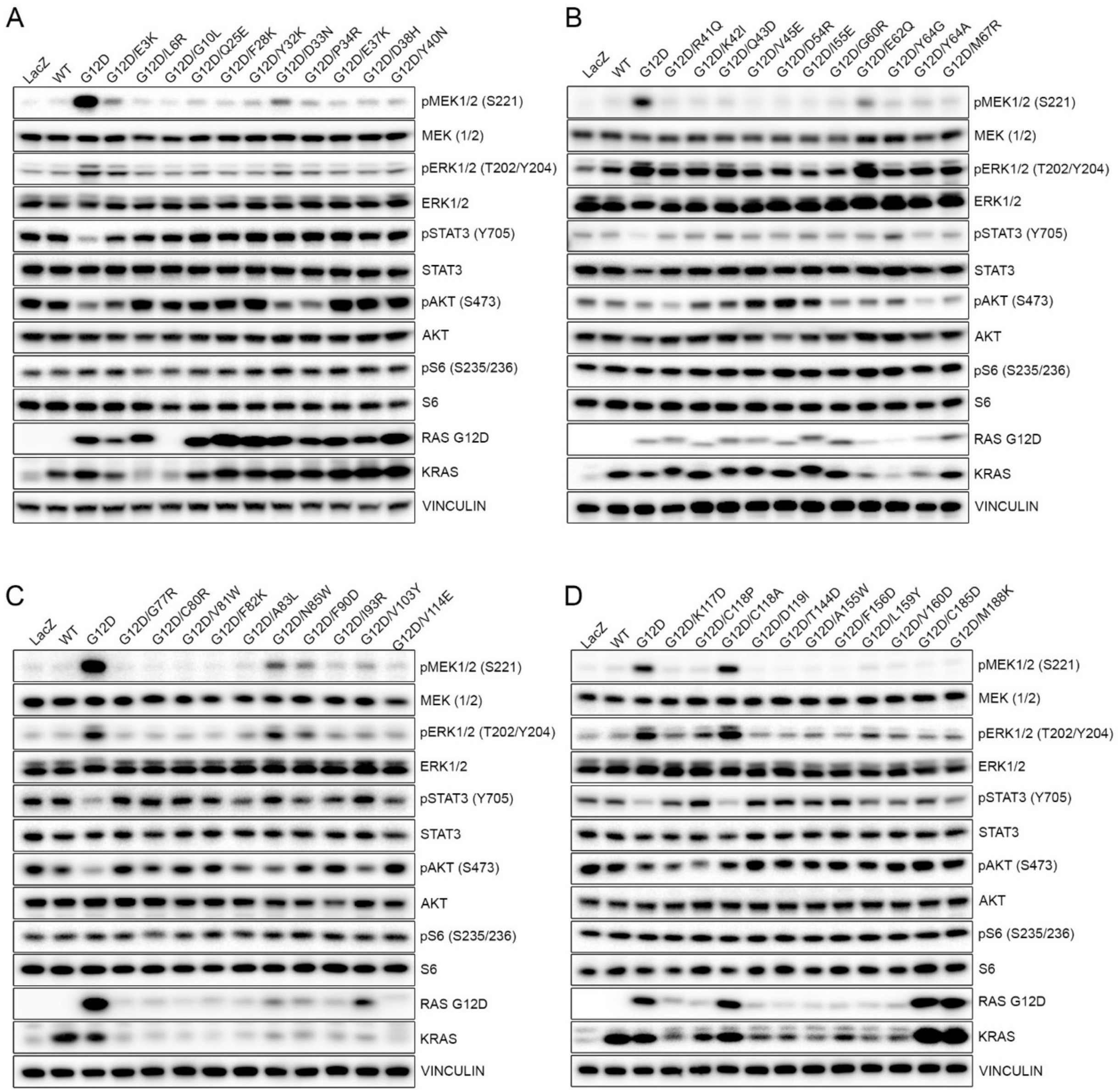


**Signaling pathways in isogenic HA1E cell lines with KRAS$^{G12D}$ alleles.** HA1E cells were transduced with lentivirus expressing indicated KRAS$^{G12D}$ alleles and positive cells were selected with puromycin. HA1E cell lines stably expressing individual allele were checked for downstream signaling with western blots.

**Extended Data Fig. 8**

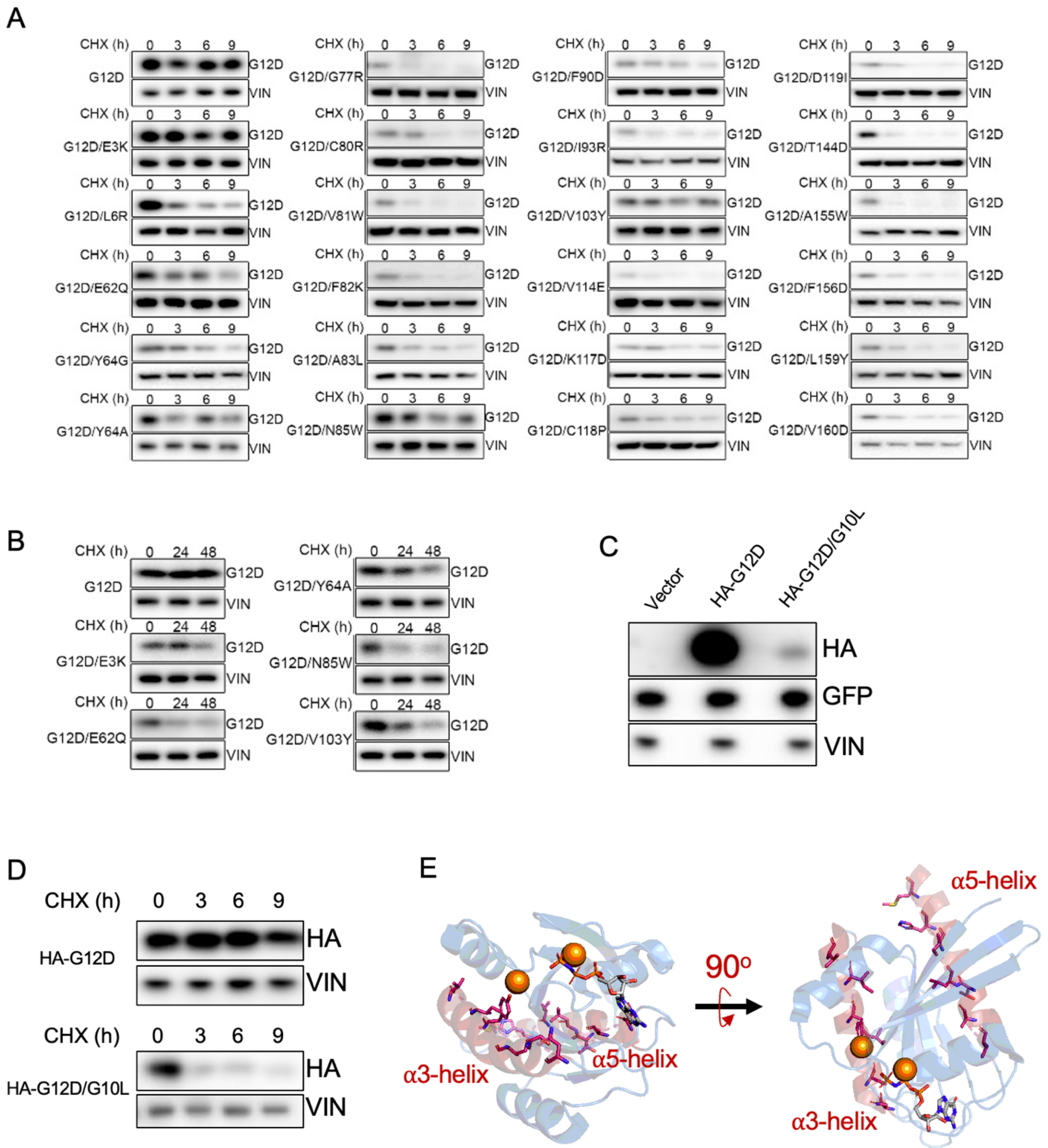


**Effect of secondary mutations on KRAS$^{G12D}$ protein stability in HA1E cells with cycloheximide (CHX) treatment.** Isogenic HA1E cell lines were treated with CHX at 20 ug/ml for 0, 3h, 6h and 9h (**A**) or for 0, 24h and 48h (**B**). KRAS$^{G12D}$ protein level was checked and vinculin was used as a loading control. (**C**) 293T cells were transfected with the vector, HA-G12D or HA-G12D/G10L and protein was harvested after 48 hours. The HA-labeled KRAS and GFP level was checked and vinculin was as a loading control. (**D**) 293T cells were transfected and cultured for 48 hours as described in (**C**). Then the cells were treated with CHX at 20 ug/ml for 0, 3h, 6h and 9h. The HA-KRAS expression was checked at indicated time points. (**E**) Structure of $KRAS^{G12D}$ bound to non-hydrolyzable GTP analog GPPNHP (PDB: 6GOF), with $\alpha3$ and $\alpha3$ helices indicated in red and side chain of residues contributing to the core of the protein are displayed.

135 **Extended Data Fig. 9**

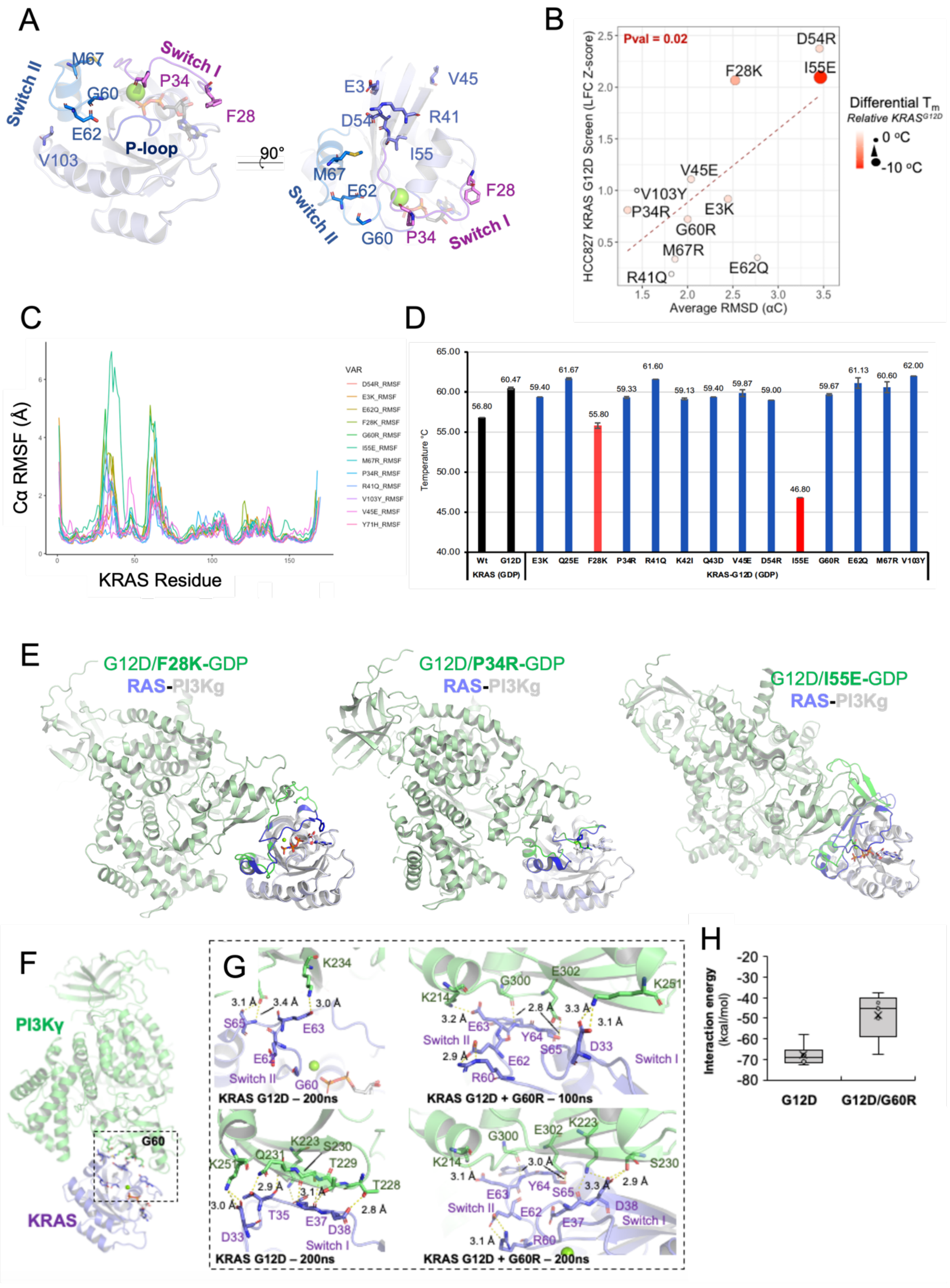


136

**Crystal structures of $KRAS^{G12D}$ inactivating mutants show conformational changes caused by mutation.** (**A**) The tertiary structure of active $KRAS^{G12D}$ (PDB: 5US4) showing the position of various $KRAS^{G12D}$ inactivating mutants that were selected for further structural and functional studies. Side-chain atoms and GDP are shown in stick representation. P-loop is colored bright blue, as well as Switch I and II regions are colored violet and cobalt, respectively. (**B**) Scatterplot of the average alpha carbon root mean square deviation (RMSD) of each $KRAS^{G12D}$ suppressor mutant crystal structure during a 200ns molecular dynamics (MD) simulation (x-axis) and the functional LFC from the $KRAS^{G12D}$ suppressor DMS screen (y-axis). Size of points indicates the differential melting temperature of each variant compared to $KRAS^{G12D}$. (**C**) The root mean square fluctuation (RMSF) of $C\alpha$ movement across a 100ns MD course for each position of $KRAS^{G12D}$ inactivating mutant structures. (**D**) Bar graph showing melting temperature (Tm) of GDP-bound WT KRAS, G12D, and G12D inactivating mutant proteins. Results are plotted as mean ± S.D ($n$ = 3). (**E**) Global view of superimposed G12D/F28K (left), G12D/P34R (middle) and G12D/I55E (right). (**F**) Global view of $KRAS^{G12D}$ modeled into HRAS-PI3Kγ complex (PDB 1HE8) with G60 residue indicated (stick representation). (**G**) Enlarged view comparing $KRAS^{G12D}$ and G12D/G60R sidechain interactions before and after 100ns MD simulation. (**H**) Box and whisker plot of KRAS-PI3Kγ interaction energy calculation predicted by Amber10 force-field-based on five representative MD simulation frames.

### Extended Data Fig. 10

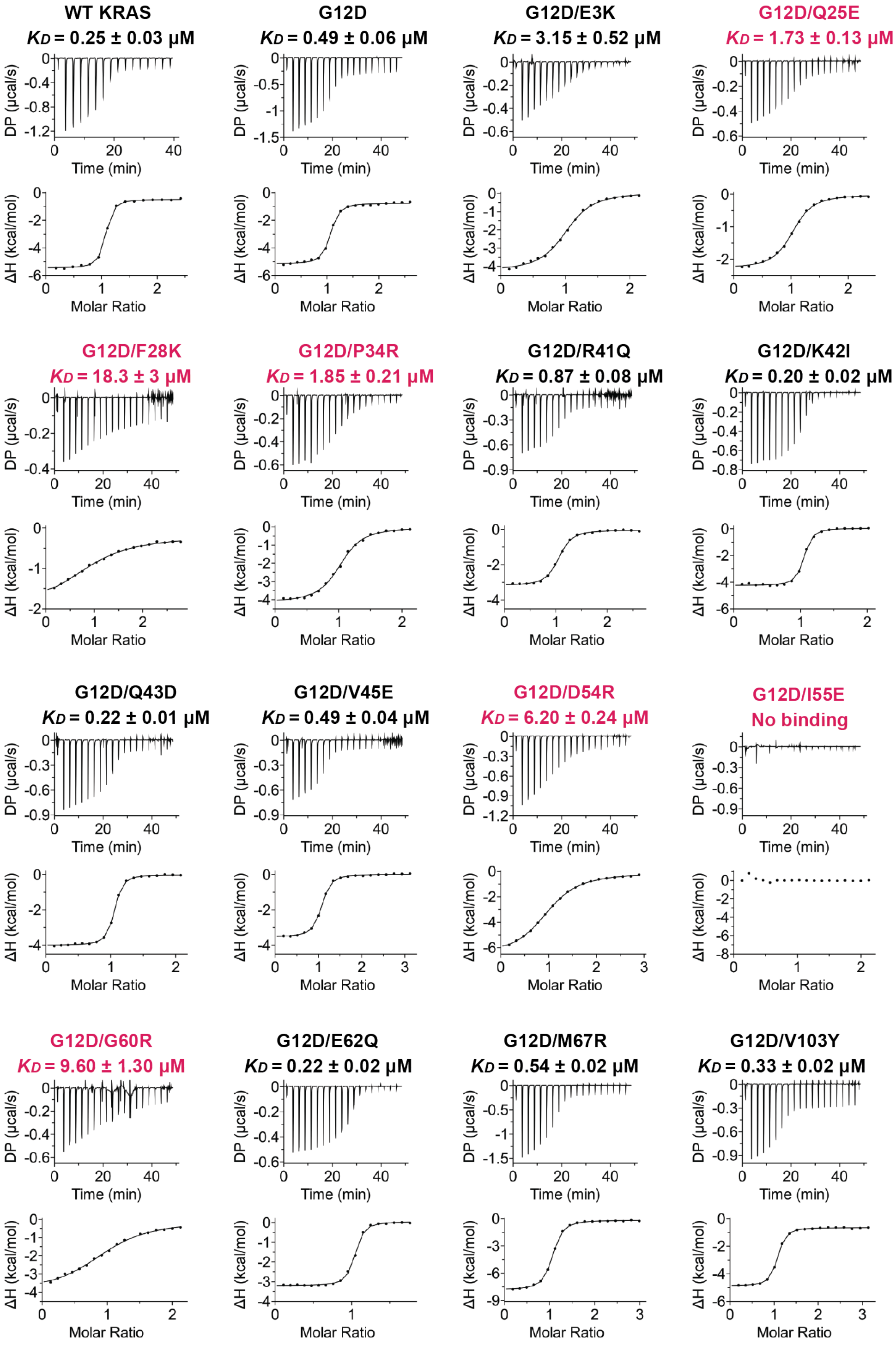


**Binding affinity measurements of KRAS$^{G12D}$ inactivating mutants with RAF1-RBD.** The dissociation constant of GMPPNP-bound WT KRAS, G12D, and G12D inactivating mutants with RAF1-RBD was measured using ITC experiments.

### Extended Data Fig. 11

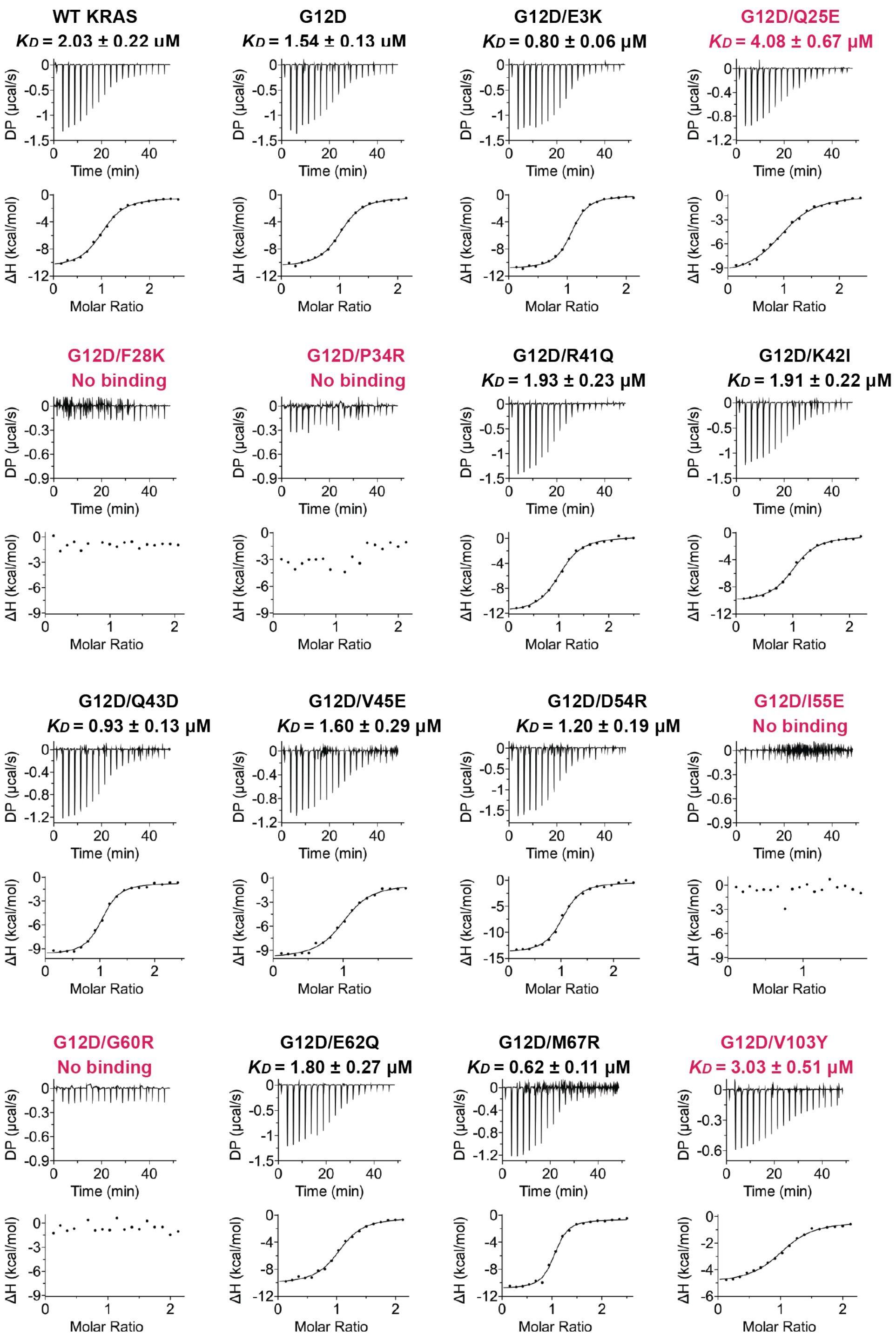


**Binding affinity measurements of KRAS$^{G12D}$ inactivating mutants with PI3Kγ.** The dissociation constant of GMPPNP-bound WT KRAS, G12D, and G12D inactivating mutants with PI3Kγ was measured using ITC experiments.

### Extended Data Fig. 12

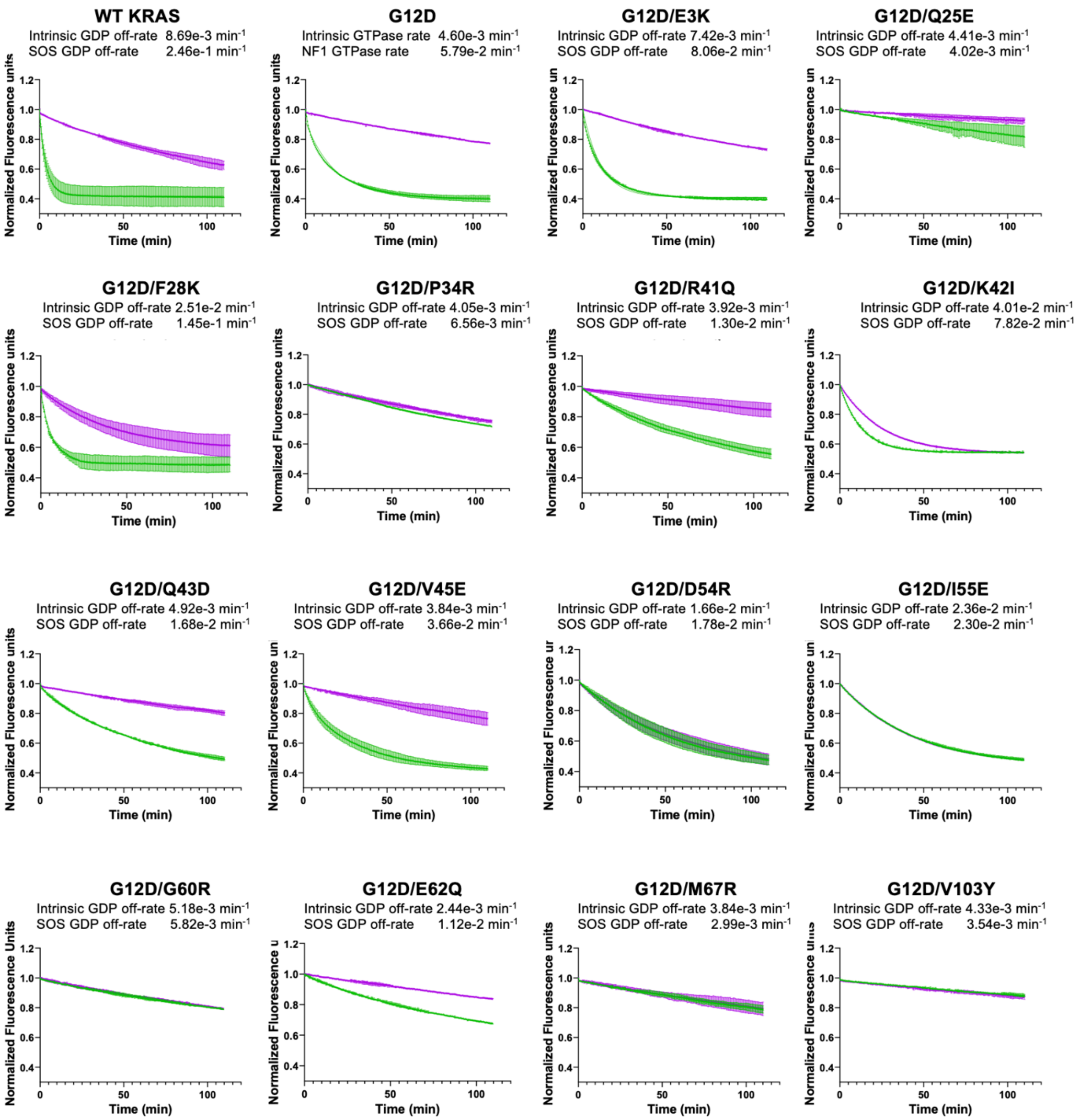


**Intrinsic and $SOS_{cat}$ mediated GDP exchange activity in $KRAS^{G12D}$ inactivating mutants**. MANT-GDP exchange from 1.5 µM was measured in the absence (purple curve) or presence (green curve) of 2.5 µM $SOS_{cat}$ in 40 mM Tris-HCl (pH 7.5), 150 mM NaCl, 2 mM $MgCl_2$, and 1 mM TCEP. The mean of replicate experiments is plotted, and error bars represent the standard deviation. Dissociation rates were calculated by fitting the data to a single exponential decay and expressed with 95% confidence intervals to show the goodness of fit.

**Extended Data Fig. 13**

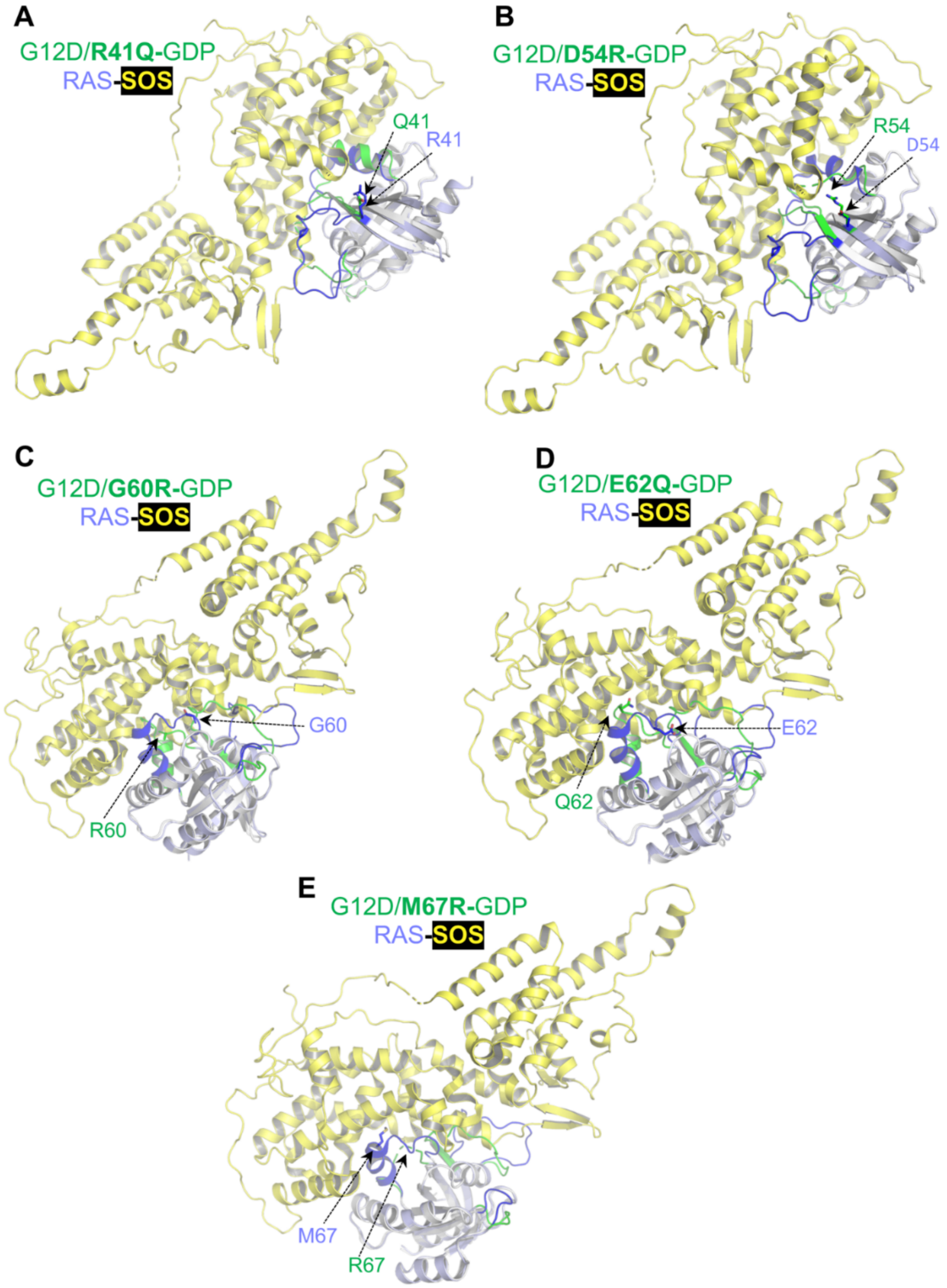


**Impact of KRAS$^{G12D}$ inactivating mutation on RAS-SOS interaction. (A-F)** Structural superposition of G12D inactivating mutants F28K (A), G60R (B), R41Q (C), Q43R (D), E62Q (E), and M67R (F) with HRAS bound at the catalytic site in the HRAS-SOS complex shows the impact of inactivating mutations on RAS-SOS interaction. SOS is colored yellow, and regions that undergo conformational changes in WT HRAS and $KRAS^{G12D}$ inactivating mutant structures are highlighted in blue and green. Side-chain atoms of the inactivating residue and GMPPNP are shown in stick representation.

**Extended Data Fig. 14**

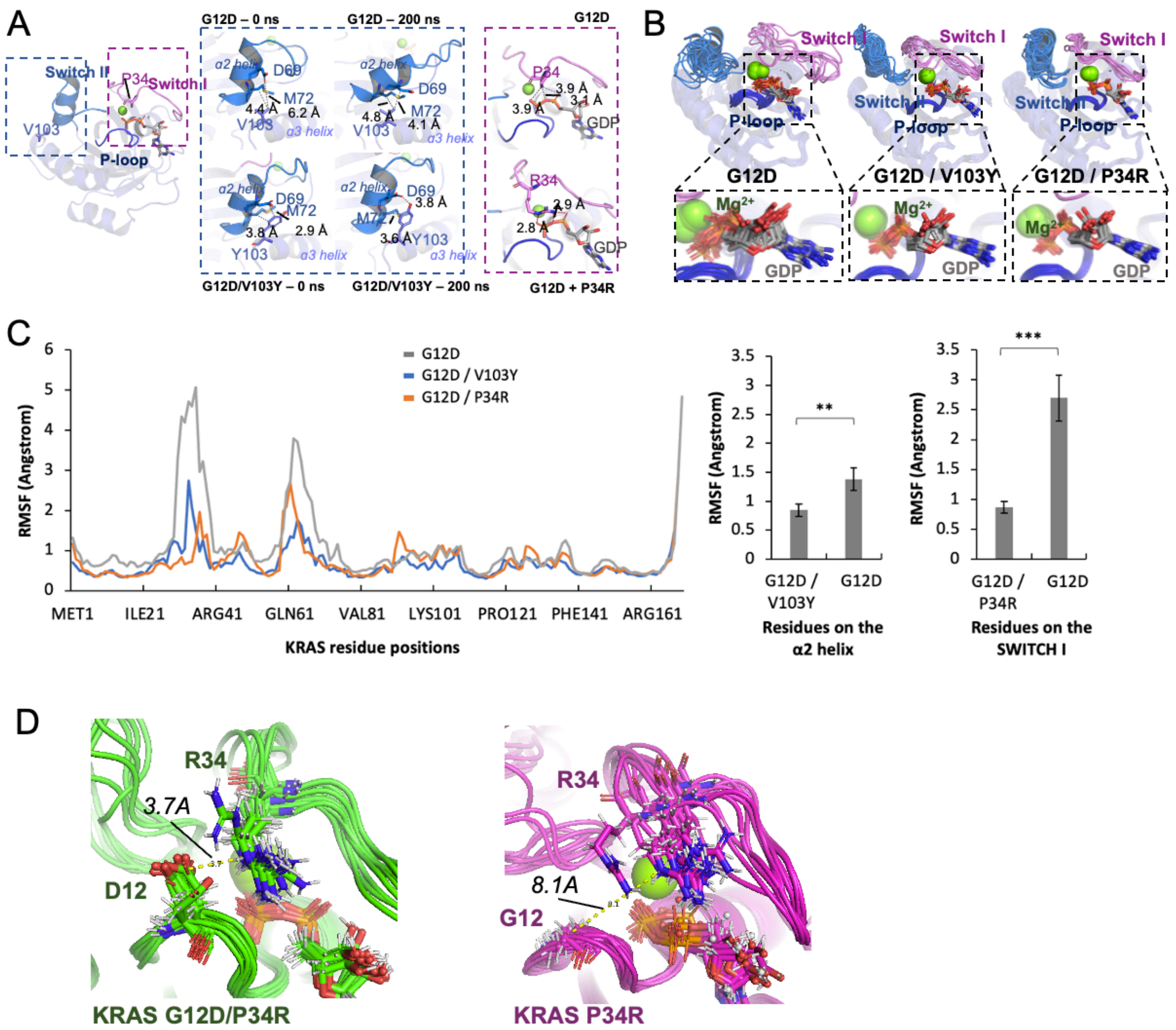


**MD simulation of KRAS$^{G12D}$ inactivating mutation and impact on GTP binding pocket.** (**A**) Global view of residues P34 and V103 indicated (stick representation) on KRAS$^{G12D}$ crystal structure (PDB 5US4). Enlarged view of structural analyses comparing KRAS$^{G12D}$ versus inactivating mutant structures G12D/V103Y and G12D/P34R. (**B**) Superposition of 100ns MD simulation frames for KRAS$^{G12D}$, G12D/V103Y, and G12D/P34R, with inset of GDP. (**C**) Per-residue root-mean-square-fluctuation (RMSF) from 100 ns molecular dynamics (MD) simulations of the KRAS$^{G12D}$ (gray trace), KRAS$^{G12D/V103Y}$ (blue trace), and KRAS$^{G12D/P34R}$ (orange trace). Mean RMSF values calculated during MD simulations for residues on the α2 helix (S65–E76) are plotted for the KRAS$^{G12D/V103Y}$ and KRAS$^{G12D}$. Mean RMSF values calculated during MD simulations for residues on the switch-I (Q25-Y40) are plotted for the KRAS$^{G12D/P34R}$ and KRAS$^{G12D}$. Comparison between two groups was performed using an unpaired, two-tailed Student's t-test. Error bars represent standard errors. **p<0.01, ***p<0.001. (**D**) Superpositions of MD trajectories of 100ns simulations on KRAS$^{G12D/P34R}$ (green) and KRAS$^{P34R}$ (magenta). Proteins are shown in cartoons. Key residues and GDP are shown in sticks.

### Extended Data Fig. 15

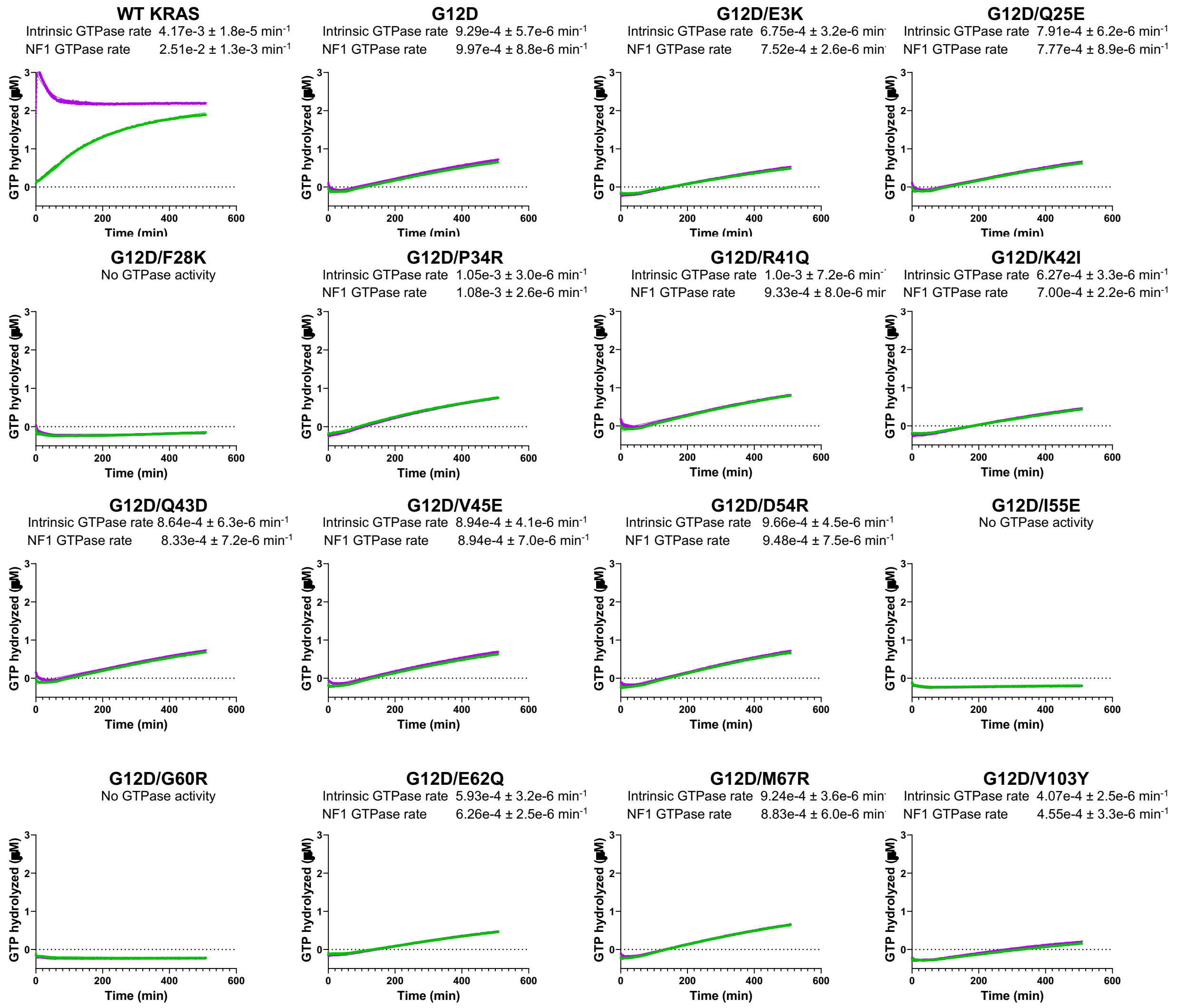


**Intrinsic and NF1 GAP mediated GTPase activity in KRAS$^{G12D}$ inactivating mutants.** GTP hydrolysis of 3 μM KRAS-GTP in the absence (green curve) or presence (purple curve) of 100 nM NF1 GAP, 50 mM Tris, pH 7.6, 150 mM NaCl, 2 mM $MgCl_2$ and 1 mM DTT was measured using the Phosphate Sensor assay. The mean of replicate experiments is plotted, and error bars represent the standard deviation. Hydrolysis rates are calculated by non-linear regression analysis and expressed with 95% confidence intervals to show the goodness of fit.

### Extended Data Fig. 16

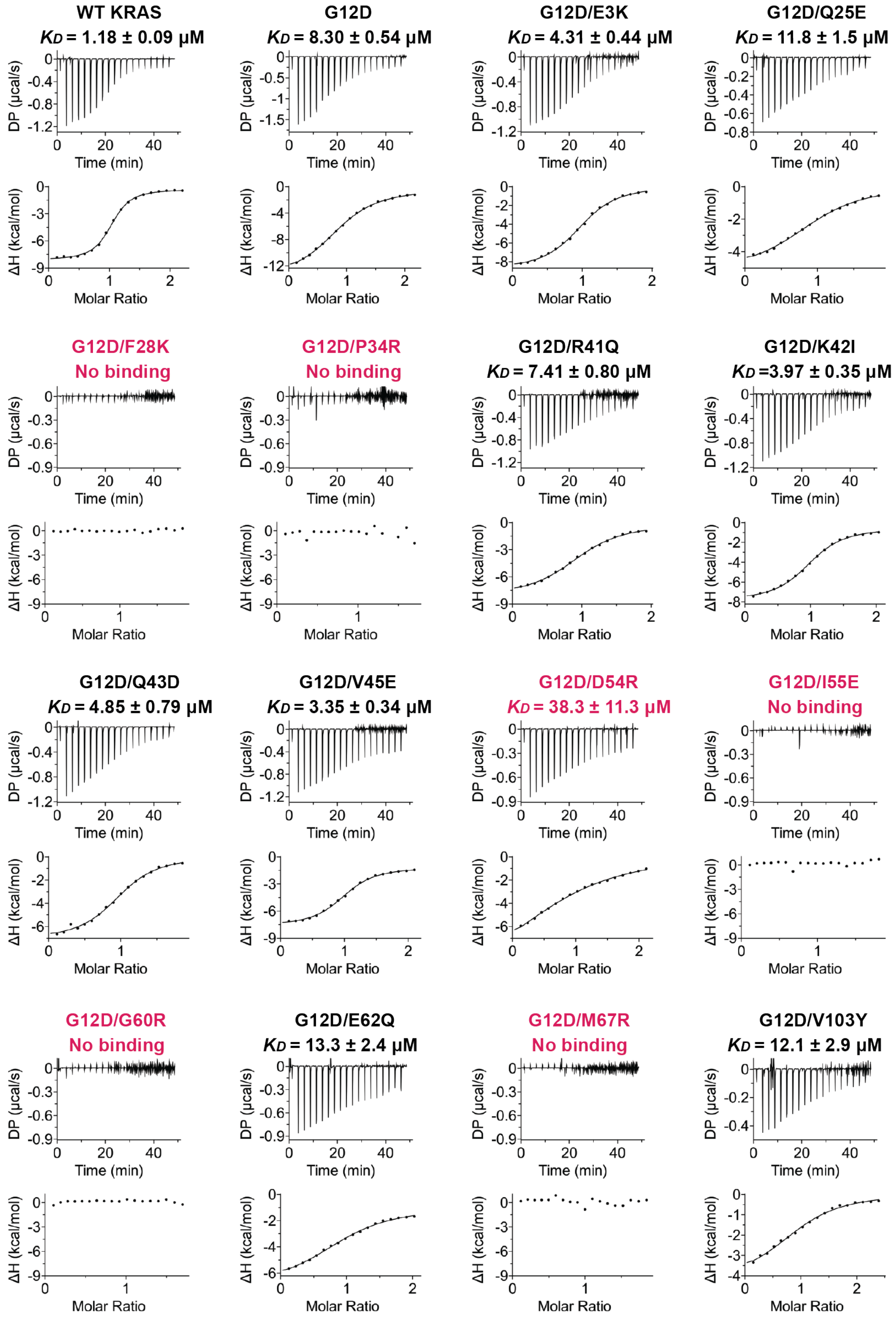


**Binding affinity measurements of KRAS$^{G12D}$ inactivating mutants with RasGAP NF1(GRD).** The dissociation constant of GMPPNP-bound WT KRAS, G12D, and G12D inactivating mutants with NF1(GRD) was measured using ITC titration experiments.
